# Radical cell identity bifurcation in *Saccharina* embryos coincides with the expression of newly acquired genes

**DOI:** 10.64898/2026.05.13.724814

**Authors:** Ioannis Theodorou, Olivier Godfroy, Samuel Boscq, Bernard Billoud, Yves Dusabyinema, Bénédicte Charrier

## Abstract

Brown algae evolved independently from animals, land plants and other algae, and we know very little about the spatio-temporal dynamics of their embryogenesis. Here, we used time-lapse, bright-field microscopy to study cell division and lineage development during early embryogenesis in the kelp *Saccharina latissima*, a large brown alga. We discovered a radical change of cell identity as early as the 4-cell stage: after fertilization, the zygote underwent two or three unequal cell divisions before the basal cell – that closest to the maternal tissue – stopped dividing and radically differentiated into a hyperpolarized cell, the rhizoid, which anchors the embryo to the substrate. RNA-seq analysis showed that differentiation of rhizoid cells was preceded by expression of 130 basal cell-specific genes. Phylostratigraphic analysis further revealed that more than 40% of these basal cell-specific genes appeared after the emergence of the brown algae group, and their functions are largely unknown. By contrast, the apical cell predominantly expressed more ancestral, metabolism-related genes, and it continued to divide to produce the long, blade-shaped thallus of the alga. The early and radical nature of cell differentiation in *Saccharina* embryos, combined with differential gene expression from various evolutionary periods, highlights the unique mechanisms of embryogenesis of this alga.

## INTRODUCTION

The process of embryogenesis differs widely among multicellular organisms across the tree of life. The rate and orientation of cell division and the dynamics of cell differentiation during embryo development vary both in time and space in different clades and species. In most animal species, the cells of the embryo divide many times before acquiring different and specific identities, although they may adopt distinct gene expression patterns as early as the two-cell stage ^1–3^. Similarly, in the embryos of land plants, the primordia of future adult tissues like the root and shoot, emerge only when the embryo is at the heart stage and comprises ∼ 500 cells ^4,5^. Brown algae are multicellular organisms that evolved independently of the evolution of animals and land plants, and even of red and green algae. Consequently, we might expect them to have different processes of embryogenesis to those of more familiar species.

Kelps are large brown algae that form dense underwater ‘forests’ in shallow temperate and Arctic oceans ^6^. An adult kelp comprises a blade-shaped lamina and stipe – which together may be up to 40 meters long – anchored to the ocean floor by a structure called the holdfast ^7,8^. Despite their curious structure and ecological and evolutionary importance, the spatio-temporal pattern of kelp embryogenesis has never been studied at the cellular and molecular levels. Here, we describe cell division and differentiation during early embryogenesis of the kelp *Saccharina latissima*. We traced cell lineage and studied spatial gene expression for up to 8 cell division rounds after the zygote stage. We found that *Saccharina* selected an original strategy of early embryogenesis, which is characterized by radical cell differentiation and the expression of species-specific genes.

## RESULTS

### Cell lineage analysis shows the basal cell abruptly ceases dividing early in embryogenesis

To identify the mechanisms involved in the embryogenesis of a kelp, we used bright field microscopy to monitor the growth of 12 embryos of the species *Saccharina latissima* (order Laminariales, family Arthrothamnaceae; ^9^) at regular intervals for up to 10 days from the zygote to the 170-cell stage (Table S1). We drew the contours of the cells of the embryos manually at each time point up to the ∼128-cell stage (Fig. S1) and inferred cell lineage by documenting each cell division event. On average, the 88-cell stage was reached after seven rounds of cell division and 192 hours (Fig. 1A; cell division is not synchronous among embryos or within a single embryo). During Phase I (1 to 8 cells; ^10^), the embryo divides only transversely, whereas during Phase II (9 to ∼1000 cells), it divides both transversely and longitudinally (Fig. 1B, top). Transverse divisions produce basal daughter cells (cells 1, 11, 12, 110, etc.) and apical daughter cells (cells 2, 21, 22, 211, etc.; see the Methods section for a detailed description of cell naming). Longitudinal divisions produce cells located on the left (A) and right (B) sides of the lamina (see Fig. 1B, top).

**Figure 1.**
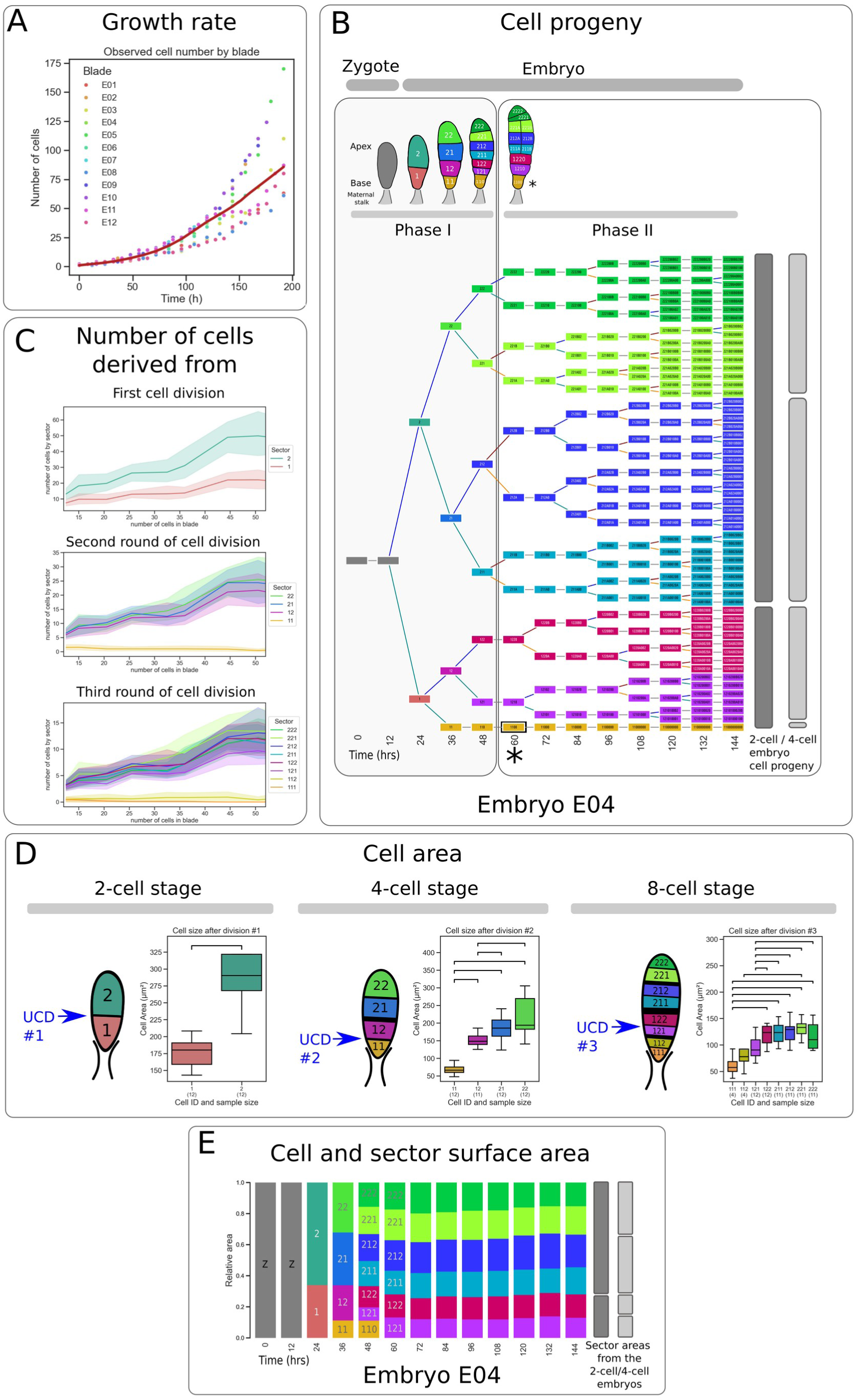
Cell lineage analysis shows the basal cell abruptly ceases dividing early in embryogenesis. **A.** Growth curve showing the number of cells per embryo for 12 embryos (E01–E12). Each dot shows one time point in time-lapse observations. The red line represents the smoothed average. **B.** Cell lineage tree of a typical embryo, E04 (see Fig. S2 for the lineage of all 12 embryos). (Top) Segmentation of the embryo is illustrated for the first 5 time points. Cells at the base of the embryo contact the maternal tissue. Cold colors (blues and greens) indicate daughters of the first apical cell in the 2-cell embryo, and warm colors (red, purple, yellow and orange) indicate daughters of the first basal cell. Cell numbers beginning with 1 are daughter cells arising from transverse cell divisions of the basal cell, and those beginning with 2 arise from similar divisions of the apical cell. The letters A and B indicate daughter cells arising from longitudinal cell divisions located on the left- and right-hand sides, respectively. Transverse cell divisions are displayed by a horizontal black lineand longitudinal cell division by a vertical line. The number 0 is given when the cell does not divide. The asterisk indicates the emergence of a rhizoid. (Bottom) Cell progeny over time. Cells are indicated with the same color code and name as in the top panel. Cell divisions are represented by straight lines, blue and green when transverse, brown and orange when longitudinal. Vertical bars on the right of the tree indicate the proportions of cell progeny that arise from the first apical and basal cells in the 2-cell (dark gray) and 4-cell embryos (light gray). The asterisk indicates the time of rhizoid emergence. Phases I and II (according to ^10^ are indicated by the light gray boxes that span the top and bottom panels. **C.** Contribution of the cells resulting from the first three rounds of cell divisions to the cell content of the growing embryos. We counted the number of cells (vertical axis) in the sectors descending from each of the two (top panel), four (middle panel) or eight (bottom panel) cells resulting from the first, second and third rounds of cell division, as the embryos grew (horizontal axis). The data shown are for all 12 embryos and the solid lines indicate the average. **D.** Measurement of cell area after unequal cell divisions in the early embryo. Schematics illustrate the embryos after the first (left; 2-cell stage), second (middle; 4-cell stage) and third (right; 8-cell stage) rounds of cell division. Black lines of various thicknesses represent the new cross walls separating two daughter cells (thicker lines indicate earlier divisions); cell numbers are as defined in panel (B). Unequal cell divisions (UCD) are indicated with a blue arrow. Box plots quantify the cell surface area (µm^2^; Y-axis) of each embryo cell at the corresponding stage. Boxes indicate the first quartile and bars the most extreme values observed up to 1.5 times the inter-quartile interval. The number of embryos quantified (sample size) is indicated below the cell ID label (X-axis). Statistically significant differences are indicated over the boxes, as determined by Mann–Whitney U test (p < 5×10^-2^). **E.** Quantification of cell and sector surface areas corresponding to cell progenies in embryo E04. Each column corresponds to a time point in embryo development from the zygote (0 hrs) to 144 cells. Each colored box corresponds to one cell (cell codes and colors as in panel (B). The height of each colored sector corresponds to the surface area of the cell or its progeny relative to that of the whole embryo. Only the eight cells resulting from the first three rounds of cell division are given distinct colors. After that, each sector aggregates the progeny of one of those eight cells.

Lineage trees revealed apico-basal asymmetry during early embryogenesis (Fig. 1B, bottom, showing embryo E04 as an example; see Fig. S2 for all 12 embryos). In 10 of the 12 embryos, the first apical cell (cell 2, green box) produced about two thirds of the embryo, whereas the basal cell (cell 1, orange box) contributed only one third (see Fig. S2 for lineage trees of all 12 embryos). We counted the number of cells descended from cell 1 and cell 2 in growing embryos with up to 50 cells in the lamina (Fig. 1C, top; Fig. S3, left column) and saw that the basal cell in the two-cell embryos produced many fewer cells than the apical cell did. A similar analysis of cells descended from cells 11, 12, 21 and 22 in the 4-cell embryo (Fig. 1C, middle; Fig. S3 middle column) showed the number of cells descended from basal cell 11 was much smaller than that of any other cell, including its sister cell 12. In the 8-cell embryo, basal cell 111 effectively produced no cells, while its sister sub-basal cell 112 produced very few compared to the other cells of the embryo (Fig. 1C, bottom; Fig. S3, right column). The apical and middle cells, by contrast, contributed new cells approximately every 12–24 hrs in Phase I, then approximately every 24–48 hrs in Phase II (Fig. S2). Thus, we conclude that division of the basal cell arrests abruptly in the second or third round of cell division.

### Arrest of cell division correlates with small basal cell size

To test whether the arrest of basal cell division correlates with cell size, we measured the surface area in 2D of each cell in the blade of the embryo. The first two daughter cells were unequal in size resulting from a first unequal cell division (UCD#1; Fig. 1D, left); the apical cell was, on average, 63% bigger than the basal cell (304.3µm^2^ vs 186.6µm^2^). A second unequal division of the basal cell was observed at the next cell division, producing a 4-cell embryo in which the upper cell (cell 12) was twice as big as the lower cell (cell 11; 151.4µm^2^ vs 73.9µm^2^; UCD#2; Fig. 1D, middle). A third unequal division was observed at the 8-cell stage when cell 12 divided to produce a larger apical cell than its sister basal cell (UCD#3; Fig. 1D, right). All the other cells from the apical progeny divided equally during Phase I. As a result, at the 8-cell stage, the embryo was composed of five upper cells of similar size, and three lower cells of gradually smaller size towards the maternal stalk (Fig. 1D, right). This establishes a longitudinal morphological asymmetry as early as the 8-cell embryo, due to a series of three unequal cell divisions.

In older embryos, the area of the ‘sectors’ formed by the progeny of the first 8 cells was roughly proportional to the size of these cells (Fig. 1E; see Fig. S4 for all the embryos). Therefore, the unequal cell divisions of Phase I impact the development of Phase II embryos in similar proportion as in Phase I embryos, exception for the most basal cell 11, which does not contribute to the growth of the lamina after the 13-cell stage, on average (Fig. 1E, orange box; Figs. S2 and S4.).

### Basal cells differentiate into rhizoids

We observed the basal cell begin to differentiate into a rhizoid in embryos comprising 5–20 cells (Fig. 2A; Table S1; Fig. S2, except E01, in which we observed the first rhizoid only at the 50-cell stage). The rhizoid is not a daughter cell of the basal cell but results from the complete and irreversible transformation of the cuboid basal cell into an elongated cell (Fig. 2B,C; Fig. S5) ^11^. The basal cell became highly polarized, it grew from the tip (Movie 1) and had few chloroplasts, whereas the cells of the lamina were cuboid and rich in chloroplasts (Fig. 2D). We observed no cell division in rhizoids, whereas the cuboid cells divided transversely and longitudinally at a regular rate (Fig. S2). The role of the rhizoid is to anchor the young embryo to the substrate ^12,13^. Throughout the embryo’s development, rhizoids are continuously produced from basal cuboid cells (Fig. 2E; Movie 1).

**Figure 2.**
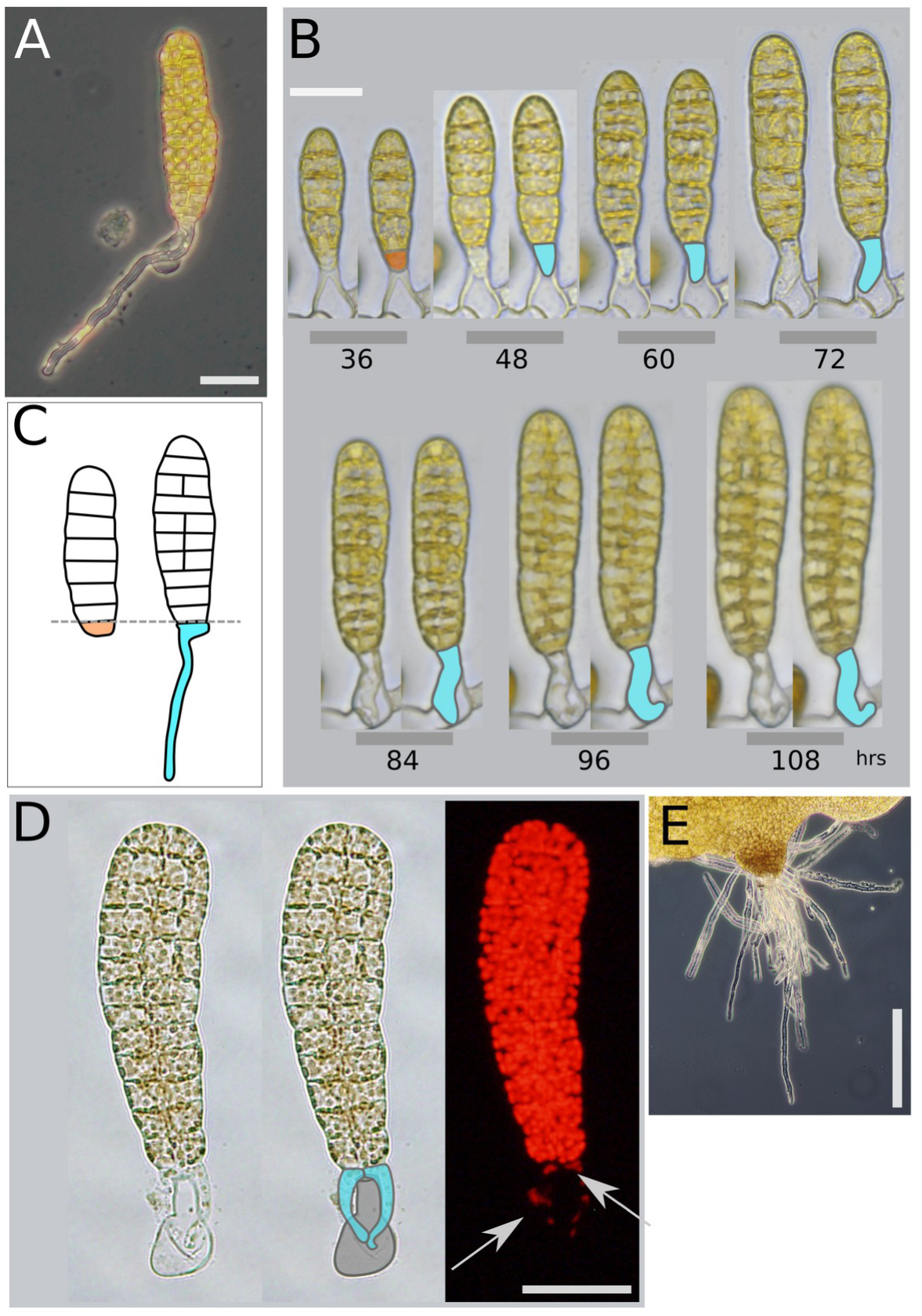
Differentiation of rhizoids from basal cells. **A.** Bright-field microscopy showing a rhizoid differentiating from the basal cell in a Phase II embryo. **B.** Time series showing differentiation of the basal cell (orange) into a rhizoid (blue) as the embryo (E02) lamina grows. **C.** Schematic summarizing differentiation of the basal cell (orange) into a rhizoid (blue). **D.** Chloroplast autofluorescence (red) in cells of the lamina and in the rhizoids (blue). The arrows indicate the few chloroplasts in the rhizoids. **E.** Bright-field microscopy showing multiple rhizoids growing from basal cells in a late embryo. Scale bars, 50 µm (A, B, D) and 100 µm (E).

The emergence of this new cell type indicates a radical bifurcation of cell identity. Basal cuboid begin to grow as an elongated rhizoid approximately 76 hours (three days) after zygote formation and 33 hours after either the second or third unequal cell division (Table S1; Figs. S2 and S4).

### Gene expression in basal cells begins to diverge in the two-cell embryo

To investigate whether the radical change in cell identity observed in the basal cell of the 13-cell embryos is accompanied or preceded by a change in gene expression, we used laser capture microdissection – a spatial transcriptomics technique enabling resolution at the single cell level ^14–17^. We captured cells and tissue sectors at various developmental stages: zygotes, apical and basal cells of 2-cell and 4-cell stage embryos (Phase I), the two median cells of 4-cell stage embryos, and the apical, median and basal sectors of Phase II embryos that contain 20, 80 and 1000 cells (∼4–13 days after fertilization; Fig. 3A; Fig. S6). We then used ultra-low input RNA sequencing (RNA-seq) to identify 5872 genes representing about one third (31.93%) of all annotated genes (Table S2a).

**Figure 3:**
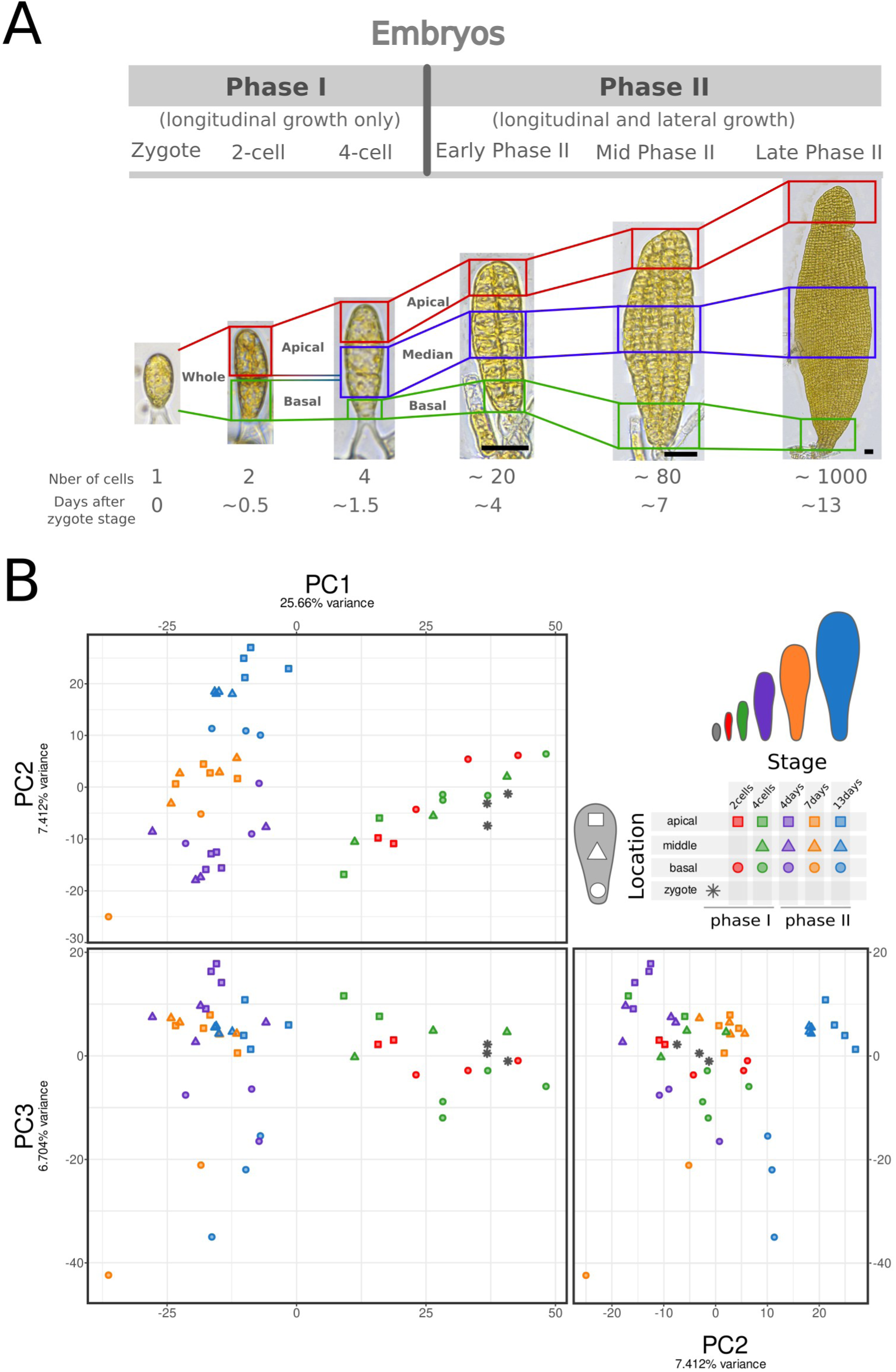
Gene expression in basal cells begins to diverge in the two-cell embryo. **A.** Laser capture microdissection of apical (red square), median (blue) and basal (green) cells and regions of embryos up to late Phase II for ultra-low input RNA sequencing. Scale bar, 30µm. n=50 Phase I embryos and n ≥ 10 of Phase II embryos were microdissected. **B.** Principal component analysis based on variance stabilized transformation (vst) of normalized counts for all analyzed genes. The first three principal components were plotted: PC1 vs PC2, upper panel; PC1 vs PC3, left lower panel left; PC2 vs PC3, right lower panel. Each point represents a replicate (n ≥3; except for apical cells of 4-cell embryos and basal cells of 7-day embryos, for which only two replicates were available). Color codes indicate developmental time points, shape codes indicate the cell or sector’s position on the longitudinal axis (see key).

Principal component analysis (PCA) based on the expression levels of those 5872 genes clustered the genes according to the embryogenesis stage (PC1 and PC2 axes; 25.66% and 7.41% variance; Fig. 3B, top left) and location along the longitudinal axis (PC2 and PC3 axes; 7.41% and 6.70% variance; Fig. 3B, bottom left and right). PC1 clearly separated Phase I and Phase II stage embryos. PC2 clustered Phase II stage embryos according to their age (4, 7 and 13 days), and separated Phase I apical cells from basal cells and zygotes. PC3 isolated the basal samples of both Phase I and Phase II. This sample grouping was confirmed by hierarchical clustering based on Spearman correlation (Fig. S7) and K.means sequential clustering (Fig. S8). Together, those three clustering methods identified marked differences between zygotes, Phase I and Phase II embryos, and a clear separation of the three Phase II embryo stages, indicating that the development stage is the most decisive parameter in determining gene expression. Superimposed on the influence of developmental stage, we found spatial organization is also an important parameter: at all stages, basal cell samples clustered away from the apical and median cell samples. This influence of spatial organization on gene expression was visible even in 2-cell stage embryos and became more pronounced as the embryos developed. Thus, the basal cell adopts a molecular identity distinct from the rest of the embryo immediately after the zygote stage.

### Gene expression reflects basal and apical cell functions

To identify the genes driving the distinct basal cell molecular identity, we used weighted gene co-expression network analysis (WGCNA), which identified 9 modules of co-expressed groups of genes (Fig. 4A). The module eigengenes, which represent the typical expression profile of the genes within the module, clustered in three main clusters that correlate either with the stage (Phase I or Phase II) or the location (apical, middle and basal) of the samples. Cluster 1 – modules brown, red, black, blue and pink – correlated positively mostly with zygote and Phase I samples. Within this cluster, modules black, blue and pink correlated strongly with Phase I, whereas modules brown and red correlated less strongly, and module black correlated strongly with the zygote stage. Cluster 2 – modules turquoise and yellow – correlated positively with Phase II embryos and apical and middle locations. Cluster 3 – modules green and gray – correlated strongly specifically with the basal cell and basal sectors. Therefore, Cluster 3 contains the genes that are involved in the differentiation of basal cells into rhizoids.

**Figure 4.**
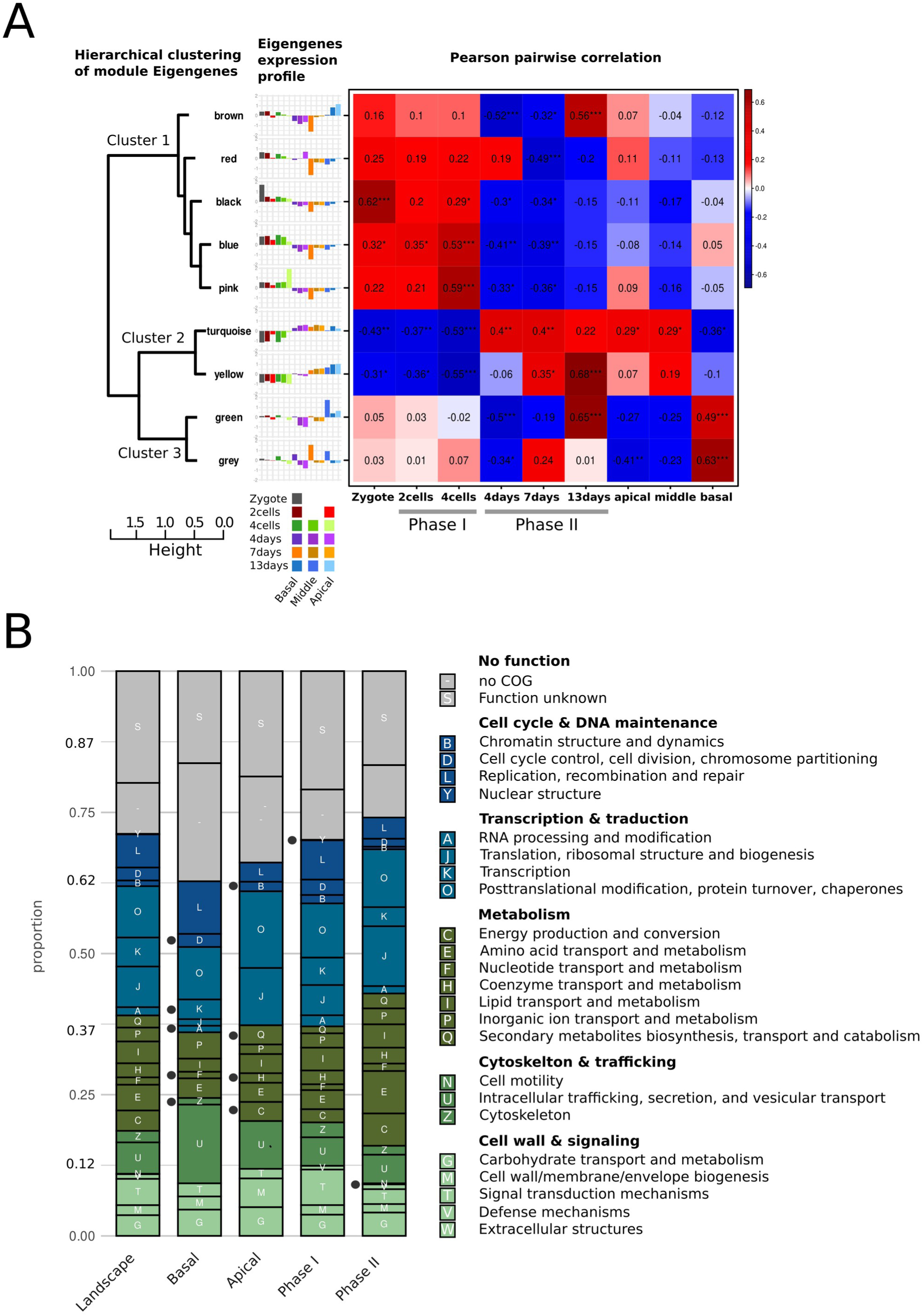
Gene expression reflects basal and apical cell functions. **A.** Weighted gene co-expression network analysis identified 9 modules of co-expressed groups of genes. (Left) Hierarchical clustering of the module eigengenes. (Middle) Bar plots of module eigengene expression: positive scores show more expression and negative scores show less expression when compared with the mean. (Right) Heat map showing pairwise Pearson correlation between module eigengene expression profile and sample stage and location. Values in the squares are Pearson correlation coefficients. Significant correlations are indicated *, P ≤0.05; **, P ≤0.01; *** P ≤0.001). **B.** Proportions of COG functional categories in the genes expressed in Basal, Apical, Phase I and Phase II groups. ‘Landscape’ indicates all genes considered in the co-expression analysis. Dots indicate COGs that are present in only one gene group.

Based on this initial gene clustering, we grouped the genes into four non-overlapping groups corresponding to Phase I, Phase II, basal and apical locations within the embryo (Fig. S9A,B). The Phase I group was defined as correlated genes from modules black, blue and pink and anti-correlated genes from modules turquoise and yellow; the Phase II group was defined as anti-correlated genes from modules black, blue and pink and correlated genes from modules turquoise and yellow; the Apical group was defined as anti-correlated genes from modules green and gray, and the Basal group was defined as correlated genes from modules green and gray (Fig. S9 A,B: Table S2a; column K).

This clustering allowed us to attribute the putative functions of every gene expressed in each group – Phase I, Phase II, Apical and Basal – by means of the Database of Clusters of Orthologous Genes (COG; https://www.ncbi.nlm.nih.gov/research/cog/<u>#;</u> Table S2a; column D). Notably, one-third of the expressed genes in the Basal group had no identified function (Fig. 4B). The remaining two-thirds included genes with putative functions in nucleotide transport and metabolism (F), transcription (K) and RNA processing (A), which were absent from the Apical group. Interestingly also, the Basal group contained genes with putative functions in control of the cell cycle (D) and in the cytoskeleton (Z), whereas the Apical group did not (Fig. 4B). The Apical group, by contrast, included genes with functions related to energy production and conversion (C), secondary metabolism, transport and catabolism (Q), and coenzymes transport and metabolism (H), which were absent from the Basal group (Fig. 4B). We conclude, therefore, that basal cells express genes that reflect the sudden arrest of their division and changes in their shape, whereas apical cells express genes related to energy and metabolite production, which reflect their involvement in the growth of the lamina. By contrast, the Phase I and Phase II groups included genes from mostly the same COG categories (Fig. 4B), although genes in the metabolism categories were more highly represented in Phase II.

To gain further insights into the potential mechanisms involved in the radical bifurcation of basal cell identity, we analyzed differential gene expression from the zygote to the 4-cell stage. The transition from the zygote to the 2-cell embryo was associated mostly with the loss of expression of some genes from both the Apical and Basal groups (Table S3a). This may reflect an early maternal-to-zygotic transition in gene expression, which was recently observed in *Undaria* ^18^, a brown alga in the order Laminariales, like *Saccharina*. Spatial differences in gene expression began to be seen in the 4-cell embryo: whereas the apical and median domains expressed similar sets of genes, at least 30 genes were differentially expressed in the basal domain. These data support those from PCA (Fig. 3), in which gene expression in the basal cell was seen to bifurcate from that of the rest of the embryo as early the 2-cell stage. During the transition from a 2-cell to a 4-cell embryo, the apical cells begin to express genes involved in the metabolism and transport of carbohydrates and other (macro)molecules (Table S3b). Starting from the 4-cell stage, the apical and median cells differ from the basal cells in their expression of genes involved in metabolic activities related to nitrogenous compounds, carbohydrates and transmembrane transport (Table S3b). This suggests an increased commitment to metabolic activities in apical and median cells as early as the 4-cell stage, which is consistent with our analysis of COG categories at all stages (Fig. 4B). By contrast, basal cells express genes involved in the biosynthesis of cell surface compounds (cell wall and cuticle) as early as the 2-cell stage. Genes that protect against oxidative stress are expressed at the 4-cell stage (Table S3b). These functions are consistent with preparing the basal cells for the emergence and growth of rhizoids.

### Genes expressed in basal cells are more recent than those in the rest of the embryo

We performed a phylostratigraphic analysis that dates the evolutionary origin of genes ^19,20^. The evolution of the transcriptomic age index (TAI), which reflects the overall age of expressed genes, shows that, during the first 13 days of embryonic development, cells and embryos from Phase I and the 13-day stage expressed younger genes than those from the 4- and 7-day stages. (Fig. S10). This profile is consistent with a transcriptomic hourglass event occurring during the early embryogenesis of *Saccharina*, as has been observed at other embryogenesis stages in different brown algae ^21^.

Furthermore, using the four groups of genes previously defined, this phylostratigraphic analysis highlights that more than 35% of the genes specifically expressed in basal cells appeared after the emergence of the brown algae group (Phaeophyceae) 450 million years ago ^22^ (Fig. 5, bottom, browns and greens; Table S2b), and 8% of these genes are specific to the order Laminariales (greens), which includes *Saccharina*. This suggests that, despite its resemblance to the rhizoids of red algae and green algae (Nagata, 1973), the rhizoid of brown algae is an innovation of brown algae. In contrast, fewer than 7% of the genes expressed in apical cells can be traced back to the emergence of brown algae (Fig. 5, top, browns and greens; Table S2), as do the genes expressed in cells located in the median sector. Thus, basal cells differ from the rest of the embryo in the significantly greater number and proportion of genes they express that emerged specifically in brown algae.

**Figure 5:**
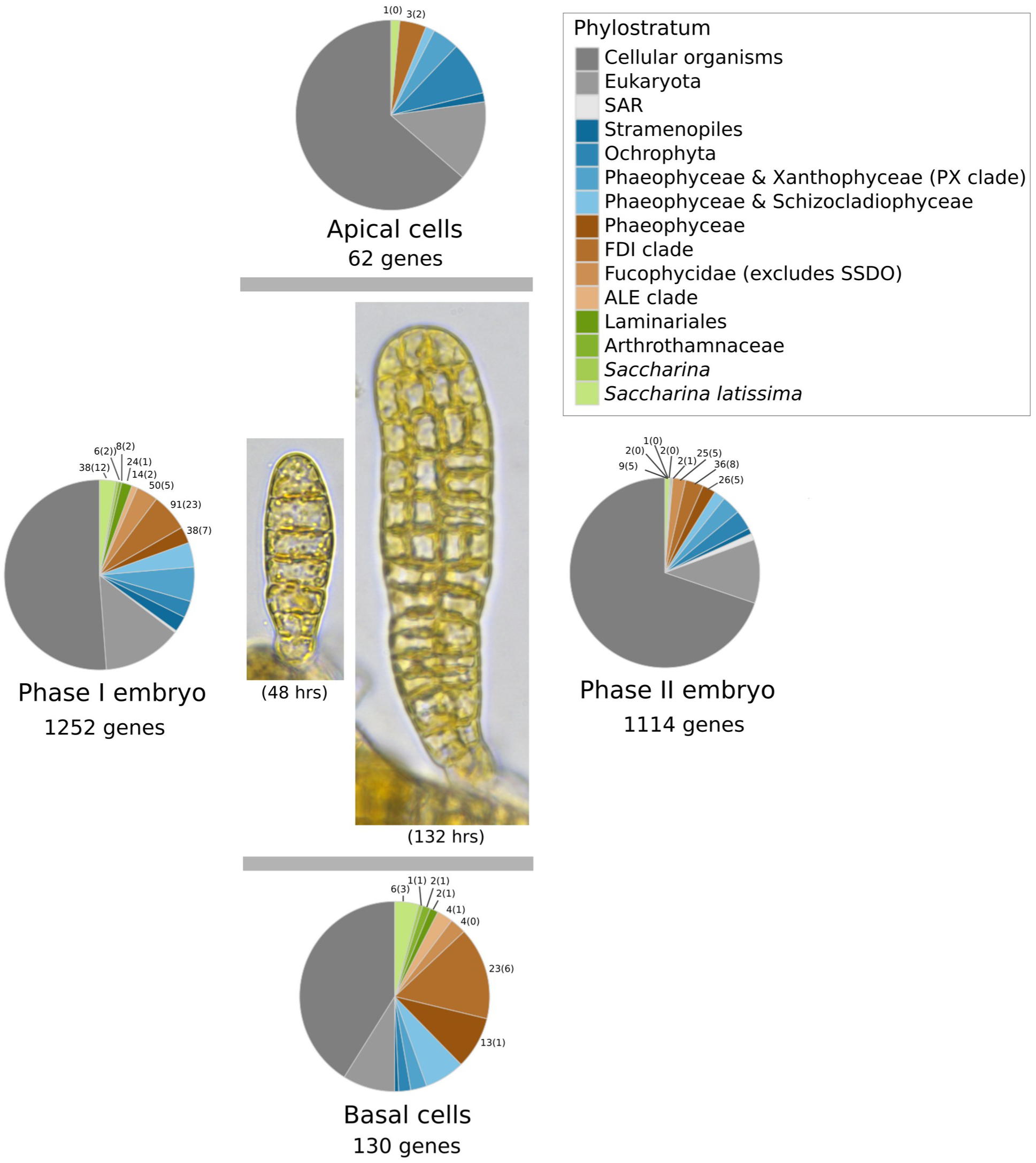
Genes expressed in basal cells are more recent than those in the rest of the embryo. For each of the four gene group preferentially expressed in Basal cells, Apical cells, Phase I or Phase II (defined in Fig. S10), the pie charts present the proportion of gene belonging to each phylostratum. Colour code indicates the position in the phylogeny. The number of genes that diverged since the emergence of brown algae (Phaeophyceae) is indicated for each phylostratum. The number of genes in each phylostratum that are similar to genes in other eukaryotes are indicated in brackets. The morphologies of Phase I and Phase II embryos, 48 and 132 hours after the zygote stage respectively, are illustrated.

This analysis also shows that Phase I embryos express a greater number of recently emerged genes than Phase II embryos. This could be explained by the hourglass pattern (Fig. S10) and by the fact that the basal region is relatively more abundant in Phase I embryos than in Phase II embryos (Fig. 5).

Of those genes that emerged specifically in brown algae, only around 25% are similar to genes with known functions. These include genes involved in response to stimuli and signaling (Tables S2b), among which are genes encoding leucine-rich repeat proteins specifically in basal cells (Table S4). Interestingly, these recent genes are also involved in intracellular transport, and RNA binding, RNA processing and RNA-mediated gene silencing, which even date back to the emergence of Laminariales (Tables S2b).

## DISCUSSION

The process by which kelp develop their embryo step by step had never been studied before. Here, we demonstrate that the fate of the basal cell in *Saccharina latissima* embryos diverges from that of the apical cell as early as the two-cell stage. This divergence is fully established after three unequal cell divisions that occur in the basal half of the embryo. As a result, by the 4-cell stage, the most basal cell stops dividing, loses its chloroplasts and by the 13-cell stage, on average, it has differentiated into a rhizoid cell. The rhizoid is the first differentiated cell type in the *Saccharina* embryo. It is an elongated, transparent cell that grows by tip growth without cell division. It appears long before the differentiation of other specialized tissues such as the meristoderm, cortex and medulla at the ∼1000-cell stage ^10^. Its emergence is preceded at the 2-cell stage by expression of 130 basal cell-specific genes. More than 40% of these genes appeared after the emergence of the brown algae group, and their functions are largely unknown. By contrast, the apical cells express predominantly more ancestral, metabolism-related genes.

The differentiation of such a specialized cell at such an early stage of embryogenesis following the first cell division is exceptional. In animals, specialized cells appear much later in development. The nematode *Caenorhabditis elegans* is described as the earliest example of cell differentiation after fertilization ^23^. Germ cell determinants named P-granules segregate in the two-cell stage embryo, but the true germ cells, Z2 and Z3, only appear at the 100-cell stage ^3,24^. In sea urchins, an unequal division at the 16-cell stage generates the micromeres, small basal blastomeres at the vegetal pole ^25,26^. Although the micromeres express specific genes already at this stage, they contribute to skeleton and germline formation only at the late gastrula stage, when the embryo contains several hundreds of cells. In mice, a recent detailed lineage study coupled with gene expression analysis precisely determined the dynamics of the emergence of fate-biased and fate-restricted cell clades before complete cell differentiation ^27^.

In plants, by contrast, cells with different morphologies and behaviors are observed as early as the 2-cell stage. In *Arabidopsis*, the first division of the elongated zygote produces the initial of the suspensor and the initial of the embryo-proper ^28^. These cells have different sizes, and patterns and dynamics of cell division. A similar process occurs in *Fucus*, a brown alga that diverged from the Laminariales 150 Mya ^22^. In this alga, the zygote elongates before the first cell division, which generates two daughter cells of different shapes and sizes, called the ‘thallus’ initial and the ‘rhizoid’ initial ^29^. However, in *Arabidopsis* as in *Fucus*, the fates of the two daughter cells are not yet distinct: in *Arabidopsis*, after several divisions of the initial suspensor cell, the uppermost cell form the quiescent center and the columella of the root tip, while the other cells of the suspensor degenerate ^30,31^. In *Fucus*, the rhizoid initial divides to form a three-dimensional tissue, which makes up about one-third of the adult thallus ^32,33^. Thus, although an initial unequal and asymmetrical cell division occurs at the zygote stage in these two plant species, the fates of their daughter cells eventually become somewhat intertwined during subsequent development ^34^. In the green alga *Ulva*, a primary rhizoid—whose morphology is more similar to that of *Saccharina* than that of *Fucus*—appears before the first division of the zygote ^35,36^. In this species, this primary rhizoid first grows by cell division along the embryo’s axis at a comparable rate to the other embryo cells ^37^. Secondary rhizoids form shortly thereafter from several stem cells derived from the thallus cell adjacent to the primary rhizoid ^36^. Therefore, in *Ulva,* rhizoids are mitotically active cells that differentiate from zygotes or stem cells. This contrasts with *Saccharina* rhizoids, which are non-dividing, hyperpolarized cells that differentiate from embryo cuboid cells, which are similar to those that grow the lamina.

Before they become rhizoids, the basal cells of *Saccharina* embryos express a high proportion of genes that recently emerged alongside brown algae and Laminariales. This is consistent with the view that the mechanisms underlying rhizoid formation in *Saccharina* embryos are, from an evolutionary perspective, distinct from those of rhizoids developing from other algae or terrestrial plants. Brown algae must anchor to the substratum as early as possible during their life cycle. Unlike most other brown algae, the zygote of *Saccharina* remains attached to the maternal tissue fixed to a substratum ^38–40^; nevertheless, the emergence of the first rhizoid reinforces the attachment of the embryo to the substratum and additional rhizoids form rapidly, which increases the survival of the organism ^12^. After about three weeks, a disk-like base tissue forms, from which the holdfast grows and supplants the attachment function of the rhizoids ^10,32,41,42^. Thus, the differentiation of the rhizoid cell identity is a transitional stage necessary for formation of a robust anchoring tissue and this step is characterized by a rapid and radical structural divergence of highly polarized rhizoid cells from the cuboid basal cells and the other cells of the thallus.

Gene expression analysis revealed that the apical and median cells are involved in metabolic functions, while the basal cells activate signaling functions. However, the exact cues that induce rhizoid differentiation have yet to be identified. The egg is a polarized cell with two flagella in the basal position. Immediately after fertilization, the intracellular organelles polarize, with vacuoles in the apical region and chloroplasts in the basal region with the maternal nucleus ^43^. Although the fused nucleus relocates to the center of the cell and the chloroplasts distribute evenly around the periphery just before the first cell division, we find that gene expression in the basal and apical cells of the 2-cell embryo differs dramatically. Therefore, it is plausible that the determinants of basal cell fate, most likely of molecular nature, originate in the egg, i.e. well before the first cell division. It is also known that in brown algae, rhizoids can differentiate in response to wounding ^44–46^. However, this seems irrelevant to differentiation after fertilization, because the base of the kelp zygote remains anchored to the stalk of the oogonium. Thus it is probably protected from physical wounding. Moreover, fertilized zygotes, and early embryos separated from the maternal stalk by sectioning, develop rhizoids at a similar developmental stage as intact embryos ^38^; Table S1). Curiously, however, parthenogenetic eggs of kelp, and zygotes that happened to be detached from the maternal stalk naturally or by shaking, develop rhizoids later than those that remain attached. Furthermore, in these embryos, the rhizoids develop in unusual positions ^11,47^. Therefore, contact between the embryo and the stalk, whether intact or sectioned, might also be an important signal for inducing the appearance of rhizoids in the correct position and at the correct time.

*Saccharina*’s response to this unknown signal is highly variable. The basal cell stops dividing between the 4-cell and 14-cell stages (on average, the 9-cell stage), and the rhizoid emerges between the 5-cell and 20-cell stages (on average, the 13-cell stage). Previous studies have reported that the early development of brown algae is morphologically plastic. For example, the spatial arrangement of elongated and round cells along the filament of the brown alga *Ectocarpus* produces up to 13 combinations at the 10-cell stage ^48^. Additionally, the lamina of early *Saccharina* embryos display length-to-width ratios ranging from 2.9 to 5.5 ^49^. In this highly variable context, at least two unequal cell divisions would be required to effectively reduce the size of the basal cell in *Saccharina* embryos and differentiate a radically different cell type, as observed in the differentiation of stomata at a more advanced stage of terrestrial plant development ^50^.

## MATERIALS AND METHODS

### Algal culture

#### F1/M1 isolation and cultivation

Because brown algae display a high variability in their development ^7,48^, we minimized genetic heterogeneity inherent to wild organism, by producing diploid embryos from two clonal parental strains, F1 (female) and M1 (male) strains, which were previously isolated in ^38^. Crushed gametophytes were diluted in Provasoli-enriched natural seawater (pNSW) to the desired concentration. 1.0 polyethylene naphthalate membrane (PEN) slides (Thermo Fisher LCM0522) placed at the bottom of 140 mm Petri dishes were previously sterilized under flow-hood UV light for 15 minutes. The gametophyte suspension was dispersed onto the sterilized PEN slides. Gametogenesis was induced by exposing gametophytes to 16 μmol photons m^-2^·s^-1^ white light with a 14:10 light-dark photoperiod at 13°C. Embryos started to develop a few days after induction.

### Morphometrics

Twelve embryos after induction ^51^ were grown at 50 μmol photons m^-2^ s^-1^ light intensity and 14:10 light:dark photoperiod at 13 °C and imaged every 8 to 24 h at a bright field inverted microscope DMI8 (Leica Microsystems GmbH) equipped with the color camera DMC4500 (Leica Microsystems©) and controlled with LAS X v3.0 (Leica Microsystems ©). The Z-stacks and their projections (average, sum, standard deviation) acquired during the time lapse were processed using Fiji ^52^ and outlines of cell contours were drawn as vector graphics using inkscape ^53^. These segmented blade images were analyzed using blade_painter ^54^ to extract morphometric data. In particular, we extracted cell area. Statistic analyses were conducted using the Mann-Whitney U test ^55^ implemented the python3 computation library scipy ^56^. Charts were drawn using the Matplotlib ^57^ and Seaborn ^58^ libraries.

In order to trace the cell lineage in embryos, we developed a suitable naming scheme, based on three rules: (1) A cell name is made of characters in {0,1,2,A,B}. The initial cell is named with an empty string. (2) For each time point, in case of division, the two daughter cells are named with the name of their common ancestor, plus one character which distinguishes them: for a transversal division, a 1 is added to the name of the most basal cell, and a 2 is added to the name of the most apical cell; for a longitudinal division, a A is added to the name of the left cell, and a B is added to the name of the right cell. If a cell does not divides, its name is elongated with a 0. So, all cells in a blade at a given time point have names with the same length. (3) Rhizoid cells are treated as not-dividing cells, and discarded from the study.

### Laser capture microdissection of *Saccharina* embryos

PEN slides with embryos growing on them were removed from pNSW and briefly rinsed (1 s) in Millipore-filtered water to eliminate salts. Slide edges were dried with tissue, and slides were immersed in 100% acetone for 30 minutes. Slides were then air-dried completely and stored at room temperature in a sealed, super-dry environment with desiccant silicate salts until laser capture microdissection. Laser micro-dissections of Phase I embryos was performed with the Carl Zeiss PALM MicroBeam system (ZEISS Axiobserver PALM DIC fluo). Embryos fixed on PEN slides were found at 5X magnification (Zeiss Fluar 5x/0.25/#420130-9900) and identified at 63X magnification (Zeiss LD Plan-NEOFLUAR 63X/0.75 Korr/#421380-9970) as they were carefully selected based on cell count and attachment to the maternal gametophyte. Using PalmRobo 4.5 software, the laser beam was guided around cell walls of cells of interest from the 1-, 2-, or 4-cell stage embryos. Laser settings were optimized to minimize radiation energy and beam diameter while ensuring effective dissection (laser energy, focus; LPC delta +; LPC focus; cutting speed; 1 cycle). Dissected cells (apical, basal, or median cells) from corresponding embryo stages were catapulted into adhesive polymer tube caps (CapSure LCM MicroCaps, ThermoScientific) using the RoboLPC function (light defocused 20 μm beneath the section). Biological replicates of 50 cells per cell-type were collected for each cell stage, with four technical replicates per type. Captures were performed over several days in a random order. Laser micro-dissections of Phase II embryos was performed using the Arcturus XT Microscope System (ThermoScientific) according to instructions from the manufacturer, with CapSure LCM MicroCaps for the captured regions. Three regions of interest (apex, middle and base) were isolated for three different stages of phase II: early phase II (5 days old, ∼20-50 cells), mid-phase II (8 days old, 100–300 cells), and late phase II (14 days old, 1500–3000 cells). The apical and basal regions comprised one fifth of the total tissue at each end, while the middle region was located between them, leaving an additional one fifth of tissue as a borderline. The captured section on top of the MicroCap was cleaned with an anatomical needle under a stereomicroscope to remove any foreign objects. Precautions were taken to minimize the sample contamination. Five biological replicates were collected per region and developmental stage. Cell numbers per replicate varied based on the region and stage; an apical region replicate from early phase II could contain around 40–50 cells, while a middle region replicate from early phase III could have between 750–1260 cells. The intensity and speed of the UV laser were optimized for each sample type to minimize tissue damage. The section obtained in MicroCaps were stored at room temperature under super-dry conditions with silica desiccants until RNA extraction.

### RNA extraction

RNA extraction was performed using the PicoPure RNA Extraction Kit (Life Technologies, #KIT0204). Immediately after laser capture microdissection, 20 µL of extraction buffer were added to the polymer tube caps. Tubes were inverted multiple times and stored at −20°C until transfer for further laboratory processing. Upon processing, extracts were reheated at 42°C for 31 minutes and centrifuged at 800 × g for 2 minutes. RNA purification was carried out following the manufacturer’s protocol. On-column DNA digestion was performed using Qiagen RNase-Free DNase I (Qiagen, #79254). RNA was eluted in 11 μL of elution buffer. Reverse transcription and amplification of the extracted RNA were conducted using the NEB Low Input RNA Amplification Kit (NewEngland BioLabs, #E6421L). Phase I samples, characterized by lower RNA yields, were amplified with 22 PCR cycles, while Phase II samples were amplified with 16 to 18 cycles. After amplification, samples were stored at −80°C. The cDNA quantity was determined using a Qubit 4 fluorometer (Thermo Fisher) and a 2100 BioAnalyzer (Agilent Technologies) with the High Sensitivity DNA Kit (Agilent Technologies, #5067-4626).

### Genomic DNA extraction

0.65 and 0.70g of fresh *Saccharina latissima* female F1 and male M1 gametophyte material, respectively, was extracted using the OmniPrep^TM^ for Plant kit (G-Biosciences, USA). Manufacturer protocol was applied with minor modifications of the lysis step: an excess of 10ml of buffer was used and complemented with 5% w/v PVP40. Genomic DNA concentration was assessed with NanoDrop One^TM^ (ThermoFisher, USA) and Qubit 4 (ThermoFisher, USA) devices and gDNA integrity was evaluated on agarose gel electrophoresis.

### gDNA and cDNA sequencing

Short read sequencing was carried out by BGI Genomics (Hong-Kong, China) using DNBSEQ™ sequencing technology, in paired-end 100bp for cDNA and 150bp for gDNA. Long-read Nanopore sequencing of the genomic DNAs of F1 and M1 strains was carried out at the IGFL Sequencing Platform (PSI). Sequencing libraries were prepared using the Ligation Sequencing Kit V14 (SQK-LSK114, ONT) and sequenced using R10.4.1 flow cells (FLO-PRO114, ONT) on a PromethION P2 solo device (ONT) with MinKNOW v23.07.12 and super-accuracy basecalling mode selected (Guppy version 7.1.4). 3.9M and 17.7M of >Q10 reads were produced respectively for M1 and F1 samples.

### Genome assembly and annotation

The genome of the M1 strain haplotype was *de novo* assembled and annotated to generate a complete reference genomic sequence with similar congruity and completeness as the other *Saccharina* genomes previously reported (Table S5 and Fig. S11).

Worklow for genome assembly and annotation is summarized in the Data Availability folder. DNBSEQ™ short reads were cleaned for low quality and remaining adapters sequence using Trimmomatic (v0.39, ^59^ and reads belonging to prokaryotic contaminants were identified and sorted out using blobtools (v1.1.1, ^60^ after a rapid meta-genomic assembly using megahit (v1.2.9, ^61^). Nanopore long reads quality was evaluated with NanoPlot (v1.42.0, ^62^. Genome assembly was carried out with NextDenovo assembler (v2.5.2, ^63^ and then polished and corrected for sequencing error with clean eukaryotic DNBSEQ™ short reads using Racon (v1.4.20, ^64^ and Pilon (v1.23, ^65^ respectively. Genome assembly congruity and completeness were then evaluated using Quast (v5.0.2, ^66^ and BUSCO (v5.5.0, ^67^ according to the eukaryota and stramenopile odb10 gene sets reference. Prior to gene structure annotation, repetitive and low complexity sequences in the genome were masked using RepeatModeler (v2.0.3, ^68^ and RepeatMasker tools (v4.1.2 – http://www.repeatmasker.org). Genome structural annotation was conduced according to BRAKER (v3.0.5, ^69^ based on RNA-seq data, *de novo* transcripts assemblies (generated with rnaSPAdes v3.15.5, ^70^ and predicted protein sequences of publicly available brown seaweed genomes. Among this BRAKER pipeline, multi-exonique genes present in the augustus.hints.gtf annotation was identified as the best *Saccharina latissima* gene set. Protein sequences predicted in those genes were functionality annotated using InterProScan (v5.68-100, ^71^ and eggNOG mapper (v2.1.12, ^72^. Orthofinder (v.2.5.2, ^73^ were used to compare the newly predicted set of *S. latissima* proteins with a subset of the publicly available brown seaweed proteome.

### RNA-seq analysis

Worklow for RNA-seq analysis is summarized in Data Availability folder. DNBSEQ™ cDNA short reads were cleaned for low quality and remaining adapters sequence using Trimmomatic (v0.39, ^59^ prior to be mapped onto the newly assembled genome with hista2 (v2.2.1, ^74^. Mapping visualization using IGV (v2.15.2, ^75^ shows that cDNA from micro-dissection mostly mapped onto the 3’-UTR regions which are less well annotated than the core of the genes. Therefore, a dedicated gene annotation was generated using StringTie (v2.1.2, ^76^ in order to retrieve UTR position coordinates. StringTie gene models overlapping genes model from BRAKER genome annotation were selected using gffcompare tool (v0.12.9, Pertea and Pertea, 2020). Then gene expression level was estimated using featureCounts (v2.0.6, ^77^. Transcriptomes profiles were then analysed on R v4.4.1. Raw counting per genes for all samples was de-noised using noisyr package (v1.0.0, ^78^ prior to be normalized using DESeq2 (v1.44.0, ^79^. Principal component analysis (PCA), Spearman correlation and k-means clustering were used to group samples according to their transcriptomics profiles using stats (v4.4.1) and factoextra (v1.0.7 – https://cran.r-project.org/web/packages/factoextra) packages respectively, and plot using ggplot2 (v3.5.1 – https://ggplot2.tidyverse.org). Gene co-expression module were generating based on normalized de-noised counts using WGCNA package (v1.73, ^80^ and GO-term enrichment was tested according to an over-representation analysis using enricher tool from clusterProfiler (v4.12.6, ^81^). TAU index of gene expression specificity was calculated using the “calc_tau” function from “scrattch.hicat” library (https://github.com/AllenInstitute/scrattch.hicat). Differential gene expression analysis of Phase I was conduced using DESeq2 package. Gene expression was represented by a z.score calculated, using normalized de-noised count, as the difference from the mean of all samples corrected by standard deviation.

### Phylostratigraphic analysis

Age of the predicted genes of our newly assembled *S. latissima* M1 strain was determined through phylostratigraphy analysis using GenEra (v1.4.2, ^19^, according to the procedure described in ^20^ (summary chart in Data Availability folder). Gene age distribution and transcriptome age index was explore on R (v4.4.1) using tidyverse (v2.0.0, ^82^ and myTAI (v0.9.3, ^83^ package respectively.

## Acknowledgement

We would like to thank Aude Le Bail and Sabine Chenivesse for maintaining and producing the F1 and M1 gametophytes. We thank the IGFL sequencing platform PSI for performing Nanopore long-read sequencing of the genomic DNAs. Pierre-Yves Canto of the IBPS-UMR7622 laser microdissection platform, and Bertrand Dubreucq and Nero Borrega of the IJPB’s Plant Observatory platform P0-Cyto are acknowledged for their support in the laser microdissection experiments. The IJPB benefits from the support of Saclay Plant Sciences-SPS (ANR-17-EUR-0007). We are grateful to B. De Reviers for valuable discussion about rhizoids in brown algae. This work is a contribution to the “ALTER e-GROW” ERC-Adv project [project number 101055148]. Views and opinions expressed are those of the author(s) only and do not necessarily reflect those of the European Union or the European Research Council Executive Agency. Neither the European Union nor the granting authority can be held responsible for them. For purposes of open access dissemination, the author applied for a CC-BY-NC license for this document.

## Data Availability

All DNA-seq and RNA-seq raw reads (short and long) have been made publicly accessible thought the European Nucleotide Archive (ENA) under the bio-project PRJEB89176. Draft genome assembly and the corresponding structural and functional annotations are available in a French academic data repository (https://entrepot.recherche.data.gouv.fr/dataverse/cnrs) accessible under the doi: https://doi.org/10.57745/TECXLY.

## Supplementary material

### Supplementary figures

**Figure S1. Time series showing the growth of an embryo of *Saccharina latissima*.**

**Figure S2. Cell lineage in twelve *Saccharina latissima* embryos.**

**Figure S3. Quantification of cellular lineage in the early embryo.**

**Figure S4. Cell size and cell sectors resulting from cell divisions in the *Saccharina* embryos.**

**Figure S5. Differentiation of basal cells into rhizoids observed in *Saccharina* embryos.**

**Figure S6. Time and location of the ablated tissues in Phase II embryos prior to RNA-seq.**

**Figure S7. Spearman correlation.**

**Figure S8. K-means sequential clustering.**

**Figure S9. Construction of groups of genes based on their expression profiles.**

**Figure S10. Transcriptome age analysis using phylostratigraphy**

**Figure S11: Genome sequence quality**

### Supplementary tables

**Table S1: Quantitative data on the time-lapse parameters and rhizoid formation.**

**Table S2: Functional annotation, phylostratigraphy, expression class and level.**

**Table S3: Dynamics of gene expression from the zygote to the embryo 4-cell stage.**

**Table S4: Identity of the recent genes expressed in the basal cells.**

**Table S5. Genome metrics of the newly assembled *Saccharina latissima* male strain M1, compared with others public genomes.**

### Supplementary movie

**Movie 1**. Embryo of *Saccharina latissima* growing in a microfluidic chip as described in ^84^. Observed in bright field microscopy for 100 hours (4 days). The differentiation of rhizoids from the basal cells is already started. The video shows rhizoids and thallus growth. The circle corresponds to one pillar of the microfluidics chip design.

## Notes

https://doi.org/10.57745/TECXLY

### Competing Interest Statement

The authors have declared no competing interest.

## References

1. Guignard, L. et al. Contact area–dependent cell communication and the morphological invariance of ascidian embryogenesis. Science 369, (2020).

2. Lawrence, P. A. & Levine, M. Mosaic and regulative development: two faces of one coin. Curr Biol 16, R236–239 (2006).

3. Sulston, J. E., Schierenberg, E., White, J. G. & Thomson, J. N. The embryonic cell lineage of the nematode *Caenorhabditis elegans*. Developmental Biology 100, 64–119 (1983).

4. Armenta-Medina, A. et al. Developmental and genomic architecture of plant embryogenesis: from model plant to crops. Plant Communications 2, 100136 (2021).

5. Jürgens, G. & Mayer, U. Arabidopsis thaliana. in Embryos Color Atlas of Development 1–18 (Wolfe Publishers, London, UK, 1994).

6. Schiel, D. R. & Foster, M. S. The Biology and Ecology of Giant Kelp Forests. (Univ of California Press, 2015).

7. Charrier, B., Le Bail, A. & Reviers, B. de. Plant Proteus: brown algal morphological plasticity and underlying developmental mechanisms. Trends in Plant Science 17, 468–477 (2012).

8. Diehl, N. et al. The sugar kelp Saccharina latissima I: recent advances in a changing climate. Annals of Botany 133, 183–212 (2024).

9. Theodorou, I. & Charrier, B. Brown Algae: Ectocarpus and Saccharina as Experimental Models for Developmental Biology. in Handbook of Marine Model Organisms in Experimental Biology 27–47 (CRC Press, Boca Raton, 2021).

10. Theodorou, I. & Charrier, B. The shift to 3D growth during embryogenesis of kelp species, atlas of cell division and differentiation of Saccharina latissima. Development 150, dev201519 (2023).

11. Sauvageau, C. Recherches sur les Laminaires des cotes de France. Mem. Acad. Sci. 56, 1–240 (1918).

12. Martínez, E. A. & Santelices, B. Selective mortality on haploid and diploid microscopic stages of *Lessonia nigrescens* Bory (Phaeophyta, Laminariales). Journal of Experimental Marine Biology and Ecology 229, 219–239 (1998).

13. Wang, W.-J., Wang, G.-C., Zhang, M. & Tseng, C. K. Isolation of Fucoxanthin from the Rhizoid of Laminaria japonica Aresch. Journal of Integrative Plant Biology 47, 1009–1015 (2005).

14. Espina, V. et al. Laser-capture microdissection. Nat Protoc 1, 586–603 (2006).

15. Gulati, G. S., D’Silva, J. P., Liu, Y., Wang, L. & Newman, A. M. Profiling cell identity and tissue architecture with single-cell and spatial transcriptomics. Nat Rev Mol Cell Biol 26, 11–31 (2025).

16. Saint-Marcoux, D., Billoud, B., Langdale, J. A. & Charrier, B. Laser capture microdissection in Ectocarpus siliculosus: the pathway to cell-specific transcriptomics in brown algae. Front Plant Sci 6, 54 (2015).

17. Saint-Marcoux, D. et al. Cell position is more important than cell shape or age for the acquisition of cell identity in the brown alga Ectocarpus. 2026.01.21.700896 Preprint at 10.64898/2026.01.21.700896 (2026).

18. Bogaert, K. A. et al. Fast zygotic genome activation in brown algae. 2026.04.21.719843 Preprint at 10.64898/2026.04.21.719843 (2026).

19. Barrera-Redondo, J., Lotharukpong, J. S., Drost, H.-G. & Coelho, S. M. Uncovering gene-family founder events during major evolutionary transitions in animals, plants and fungi using GenEra. Genome Biol 24, 54 (2023).

20. Denoeud, F. et al. Evolutionary genomics of the emergence of brown algae as key components of coastal ecosystems. Cell 187, 6943–6965.e39 (2024).

21. Lotharukpong, J. S. et al. A transcriptomic hourglass in brown algae. Nature 1–7 (2024) doi:10.1038/s41586-024-08059-8.

22. Choi, S.-W. et al. Ordovician origin and subsequent diversification of the brown algae. Current Biology 34, 740–754.e4 (2024).

23. Gilbert, S. F. Developmental Biology. (Sinauer Associates, Sunderland, Mass, 2000).

24. Hird, S. N., Paulsen, J. E. & Strome, S. Segregation of germ granules in living Caenorhabditis elegans embryos: cell-type-specific mechanisms for cytoplasmic localisation. Development 122, 1303–1312 (1996).

25. Emura, N. & Yajima, M. Micromere formation and its evolutionary implications in the sea urchin. Curr Top Dev Biol 146, 211–238 (2022).

26. Logan, C. Y., Miller, J. R., Ferkowicz, M. J. & McClay, D. R. Nuclear β-catenin is required to specify vegetal cell fates in the sea urchin embryo. Development 126, 345–357 (1999).

27. Colgan, W. N. et al. Comprehensive Lineage Tracing Maps the Landscape of Cell Fate Decisions in Mouse Embryogenesis. 2026.05.07.722278 Preprint at 10.64898/2026.05.07.722278 (2026).

28. Lau, S., Slane, D., Herud, O., Kong, J. & Jürgens, G. Early Embryogenesis in Flowering Plants: Setting Up the Basic Body Pattern. Annual Review of Plant Biology 63, 483–506 (2012).

29. Bouget, F. Y., Berger, F. & Brownlee, C. Position dependent control of cell fate in the Fucus embryo: role of intercellular communication. Development 125, 1999–2008 (1998).

30. Downs, J. & Jones, B. The short and intricate life of the suspensor. Physiologia Plantarum 169, 110–121 (2020).

31. Jürgens, G. Apical–basal pattern formation in Arabidopsis embryogenesis. EMBO J 20, 3609–3616 (2001).

32. Fritsch, F. E. The Structure And Reproduction Of The Algae. vol. 2 (Cambridge University Press, Cambridge, 1945).

33. Nienburg, W. Die Entwicklung der Keimlinge von Fucus vesiculosus und ihre Bedeutung für die Phylogenie der Phaeophyceen. Wissenschaftliche Meeresuntersuchungen: Abteilung Kiel 52–62 (1931).

34. Charrier, B. Asymmetrical cell division in brown algae: How far can we take the paradigm? Current Opinion in Plant Biology 86, 102758 (2025).

35. Gayral, P. Morphologie et morphogénèse des Ulvales. Soc. bot Fr., Mém. 115:sup1, 130–141 (1968).

36. Wichard, T. et al. The green seaweed Ulva: a model system to study morphogenesis. Front Plant Sci 6, 72 (2015).

37. Fjeld, A. Genetic control of cellular differentiation in *Ulva mutabilis*. Gene effects in early development. Developmental Biology 28, 326–343 (1972).

38. Boscq, S. et al. MUM, a maternal unknown message, inhibits early establishment of the medio-lateral axis in the embryo of the kelp Saccharina latissima. Development 151, dev202732 (2024).

39. Fletcher, R. L. & Callow, M. E. The settlement, attachment and establishment of marine algal spores. British Phycological Journal 27, 303–329 (1992).

40. Veenhof, R. J. et al. Kelp gametophytes in changing oceans. in Oceanography and marine biology 335–371 (CRC Press, 2022).

41. Davies, J. M., Ferrier, N. C. & Johnston, C. S. The Ultrastructure of the Meristoderm Cells of the Hapteron of *Laminaria*. J. Mar. Biol. Ass. 53, 237–246 (1973).

42. Drew, G. H. The Reproduction and early Development of *Laminaria digitata* and *Laminaria saccharina*. Annals of Botany 24, 177–189 (1910).

43. Motomura, T. Ultrastructure of Fertilization in *Laminaria angustata* (Phaeophyta, Laminariales) with Emphasis on the Behavior of Centrioles, Mitochondria and Chloroplasts of the Sperm. Journal of Phycology 26, 80–89 (1990).

44. Russell, G. Rhizoid production in excised Dictyota dichotoma. British Phycological Journal 5, 243–245 (1970).

45. Shirae-Kurabayashi, M., Edzuka, T., Suzuki, M. & Goshima, G. Cell tip growth underlies injury response of marine macroalgae. PLOS ONE 17, e0264827 (2022).

46. Tanaka, A., Hoshino, Y., Nagasato, C. & Motomura, T. Branch regeneration induced by sever damage in the brown alga Dictyota dichotoma (dictyotales, phaeophyceae). Protoplasma 254, 1341–1351 (2017).

47. Dries, E. et al. Cell wall-mediated maternal control of apical–basal patterning of the kelp Undaria pinnatifida. New Phytologist 243, 1887–1898 (2024).

48. Le Bail, A., Billoud, B., Le Panse, S., Chenivesse, S. & Charrier, B. ETOILE regulates developmental patterning in the filamentous brown alga Ectocarpus siliculosus. Plant Cell 23, 1666–1678 (2011).

49. Boscq, S. et al. Longitudinal growth of the Saccharina kelp embryo depends on actin filaments that control the formation of a corset-like structure composed of alginate. Sci Rep 15, 1178 (2025).

50. Mathew, M. M. & Bergmann, D. C. Mechanisms and flexibility in plant asymmetric cell division. Current Opinion in Plant Biology 91, 102896 (2026).

51. Theodorou, I., Opsahl-Sorteberg, H.-G. & Charrier, B. Preparation of Zygotes and Embryos of the Kelp Saccharina latissima for Cell Biology Approaches. Bio-protocol e4132–e4132 (2021).

52. Schindelin, J. et al. Fiji - an Open Source platform for biological image analysis. Nature methods 9, 676–682 (2012).

53. Inkscape. Inkscape Project. (2022).

54. Boscq, S., Billoud, B. & Charrier, B. Cell-Autonomous and Non-Cell-Autonomous Mechanisms Concomitantly Regulate the Early Developmental Pattern in the Kelp Saccharina latissima Embryo. Plants 13, 1341 (2024).

55. Mann, H. B. & Whitney, D. R. On a Test of Whether one of Two Random Variables is Stochastically Larger than the Other. The Annals of Mathematical Statistics 18, 50–60 (1947).

56. Virtanen, P. et al. SciPy 1.0: fundamental algorithms for scientific computing in Python. Nat Methods 17, 261–272 (2020).

57. Hunter, J. D. Matplotlib: A 2D Graphics Environment. Computing in Science & Engineering 9, 90–95 (2007).

58. Waskom, M. L. seaborn: statistical data visualization. The Journal of Open Source Software 6, 3021 (2021).

59. Bolger, A. M., Lohse, M. & Usadel, B. Trimmomatic: a flexible trimmer for Illumina sequence data. Bioinformatics 30, 2114–2120 (2014).

60. Laetsch, D. R. & Blaxter, M. L. BlobTools: Interrogation of genome assemblies. Preprint at 10.12688/f1000research.12232.1 (2017).

61. Li, D., Liu, C.-M., Luo, R., Sadakane, K. & Lam, T.-W. MEGAHIT: an ultra-fast single-node solution for large and complex metagenomics assembly via succinct de Bruijn graph. Bioinformatics 31, 1674–1676 (2015).

62. De Coster, W. & Rademakers, R. NanoPack2: population-scale evaluation of long-read sequencing data. Bioinformatics 39, btad311 (2023).

63. Hu, J. et al. NextDenovo: an efficient error correction and accurate assembly tool for noisy long reads. Genome Biol 25, 107 (2024).

64. Vaser, R., Sović, I., Nagarajan, N. & Šikić, M. Fast and accurate de novo genome assembly from long uncorrected reads. Genome Res. 27, 737–746 (2017).

65. Walker, B. J. et al. Pilon: An Integrated Tool for Comprehensive Microbial Variant Detection and Genome Assembly Improvement. PLOS ONE 9, e112963 (2014).

66. Gurevich, A., Saveliev, V., Vyahhi, N. & Tesler, G. QUAST: quality assessment tool for genome assemblies. Bioinformatics 29, 1072–1075 (2013).

67. Seppey, M., Manni, M. & Zdobnov, E. M. BUSCO: Assessing Genome Assembly and Annotation Completeness. in Gene Prediction: Methods and Protocols (ed. Kollmar, M.) 227–245 (Springer, New York, NY, 2019). doi:10.1007/978-1-4939-9173-0_14.

68. Flynn, J. M. et al. RepeatModeler2 for automated genomic discovery of transposable element families. Proceedings of the National Academy of Sciences 117, 9451–9457 (2020).

69. Gabriel, L. et al. BRAKER3: Fully automated genome annotation using RNA-seq and protein evidence with GeneMark-ETP, AUGUSTUS, and TSEBRA. Genome Res. 34, 769–777 (2024).

70. Bushmanova, E., Antipov, D., Lapidus, A. & Prjibelski, A. D. rnaSPAdes: a de novo transcriptome assembler and its application to RNA-Seq data. Gigascience 8, giz100 (2019).

71. Jones, P. et al. InterProScan 5: genome-scale protein function classification. Bioinformatics 30, 1236–1240 (2014).

72. Cantalapiedra, C. P., Hernández-Plaza, A., Letunic, I., Bork, P. & Huerta-Cepas, J. eggNOG-mapper v2: Functional Annotation, Orthology Assignments, and Domain Prediction at the Metagenomic Scale. Mol Biol Evol 38, 5825–5829 (2021).

73. Emms, D. M. & Kelly, S. OrthoFinder: phylogenetic orthology inference for comparative genomics. Genome Biol 20, 238 (2019).

74. Kim, D., Paggi, J. M., Park, C., Bennett, C. & Salzberg, S. L. Graph-based genome alignment and genotyping with HISAT2 and HISAT-genotype. Nat Biotechnol 37, 907–915 (2019).

75. Robinson, J. T. et al. Integrative genomics viewer. Nat Biotechnol 29, 24–26 (2011).

76. Pertea, M. et al. StringTie enables improved reconstruction of a transcriptome from RNA-seq reads. Nat Biotechnol 33, 290–295 (2015).

77. Liao, Y., Smyth, G. K. & Shi, W. featureCounts: an efficient general purpose program for assigning sequence reads to genomic features. Bioinformatics 30, 923–930 (2014).

78. Moutsopoulos, I. et al. noisyR: enhancing biological signal in sequencing datasets by characterizing random technical noise. Nucleic Acids Res 49, e83 (2021).

79. Love, M. I., Huber, W. & Anders, S. Moderated estimation of fold change and dispersion for RNA-seq data with DESeq2. Genome Biol 15, 550 (2014).

80. Zhang, B. & Horvath, S. A General Framework for Weighted Gene Co-Expression Network Analysis. Statistical Applications in Genetics and Molecular Biology 4, (2005).

81. Yu, G., Wang, L.-G., Han, Y. & He, Q.-Y. clusterProfiler: an R Package for Comparing Biological Themes Among Gene Clusters. OMICS: A Journal of Integrative Biology 16, 284–287 (2012).

82. Wickham, H. et al. Welcome to the Tidyverse. Journal of Open Source Software 4, 1686 (2019).

83. Drost, H.-G., Gabel, A., Liu, J., Quint, M. & Grosse, I. myTAI: evolutionary transcriptomics with R. Bioinformatics 34, 1589–1590 (2018).

84. Clerc, T. et al. Cultivation and Imaging of S. latissima Embryo Monolayered Cell Sheets Inside Microfluidic Devices. Bioengineering 9, 718 (2022).

