## Supplementary figures for "Radical cell identity bifurcation in *Saccharina* embryos coincides with the expression of newly acquired genes"

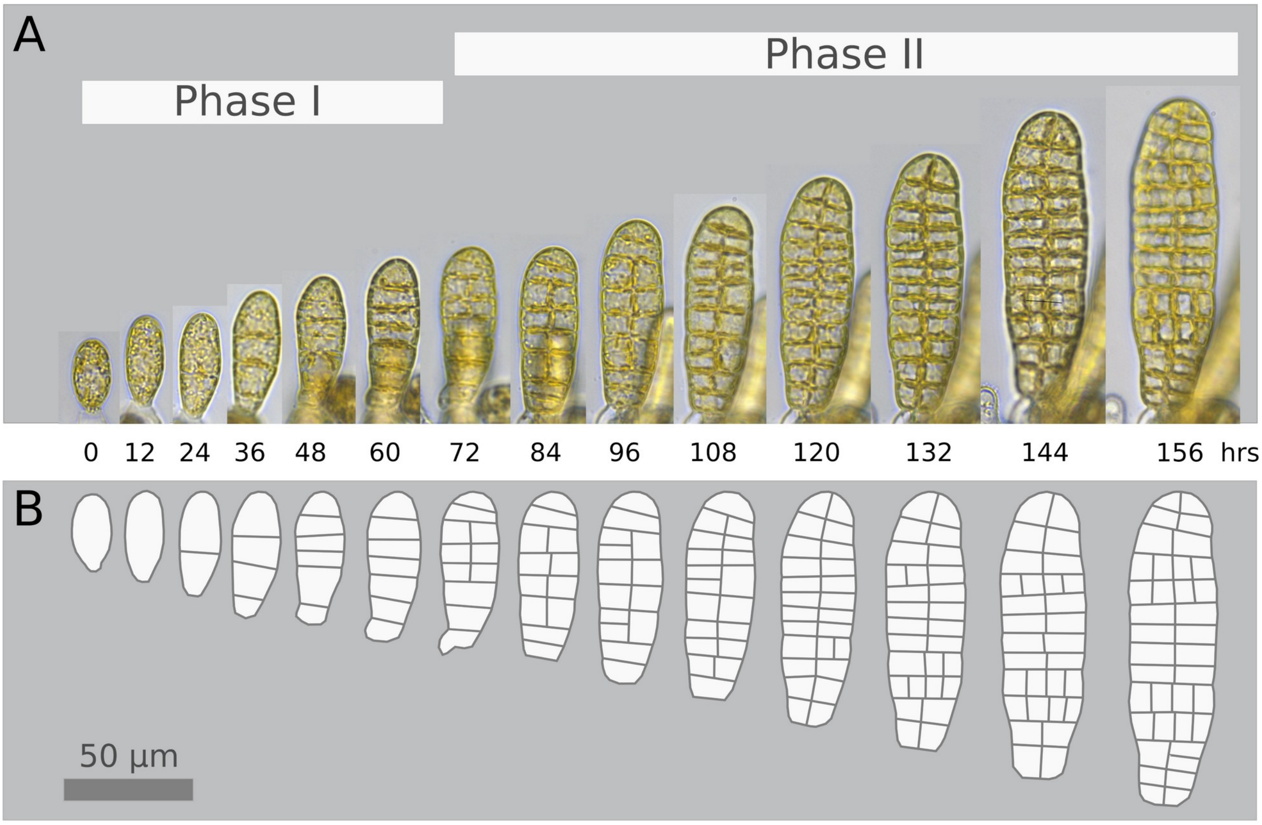


**Figure S1. Time series showing the growth of an embryo of *Saccharina latissima*.**

**A.** The time series starts at the zygote stage (0 hours) and continues for 156 hours (final time point). The embryo progresses through Phases I and II of embryogenesis. Phase I is characterized by transverse cell divisions only, and Phase II is characterized by both transverse and longitudinal cell divisions. **B.** The cell contours were drawn manually.

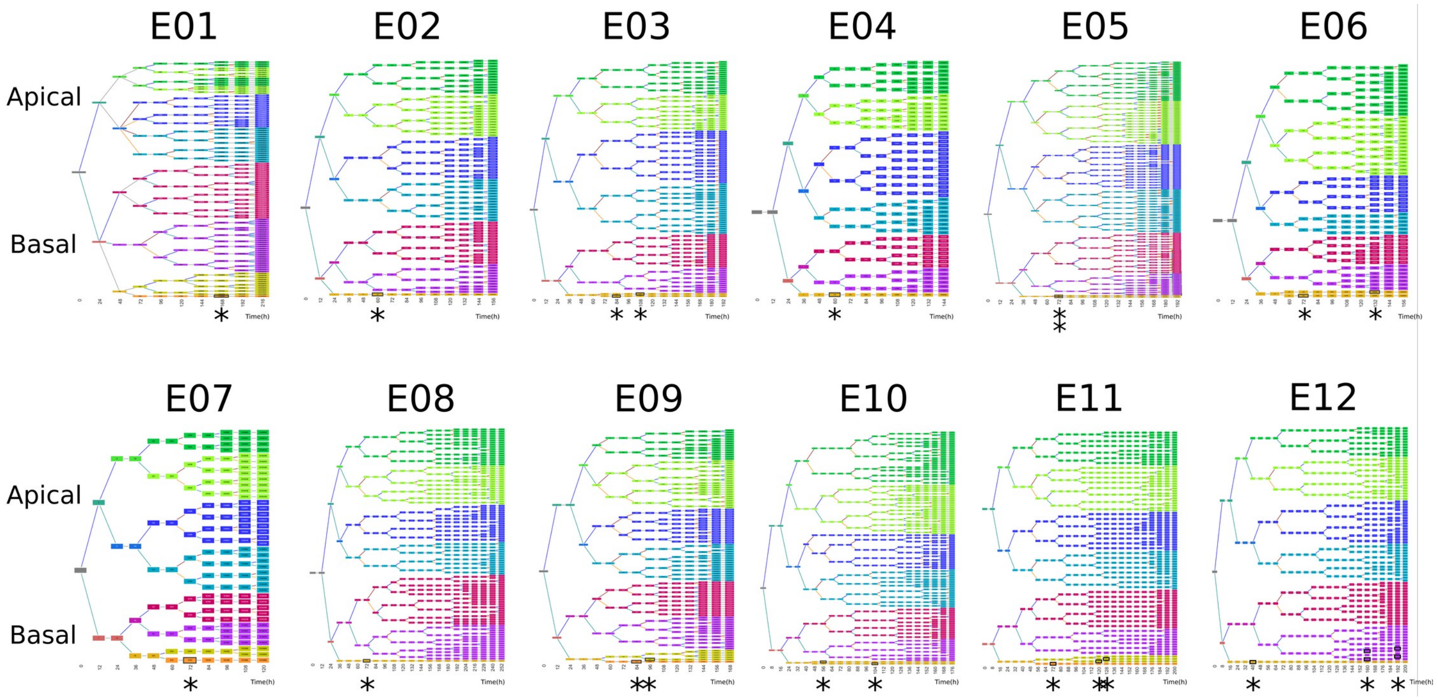


**Figure S2. Cell lineage in twelve *Saccharina latissima* embryos.**

The cell lineage tree of the 12 embryos (E01 to E12) is shown, with the same color code and legend as in Fig. 1B. Asterisks below the tree indicate the time when the basal cell differentiates into a rhizoid. Boxes with a black frames indicate cells that have differentiated into rhizoids. The positions of the apical (cool colors) and the basal (warm colors) cells of the 2-cell stage embryo are indicated on the left side of the tree.

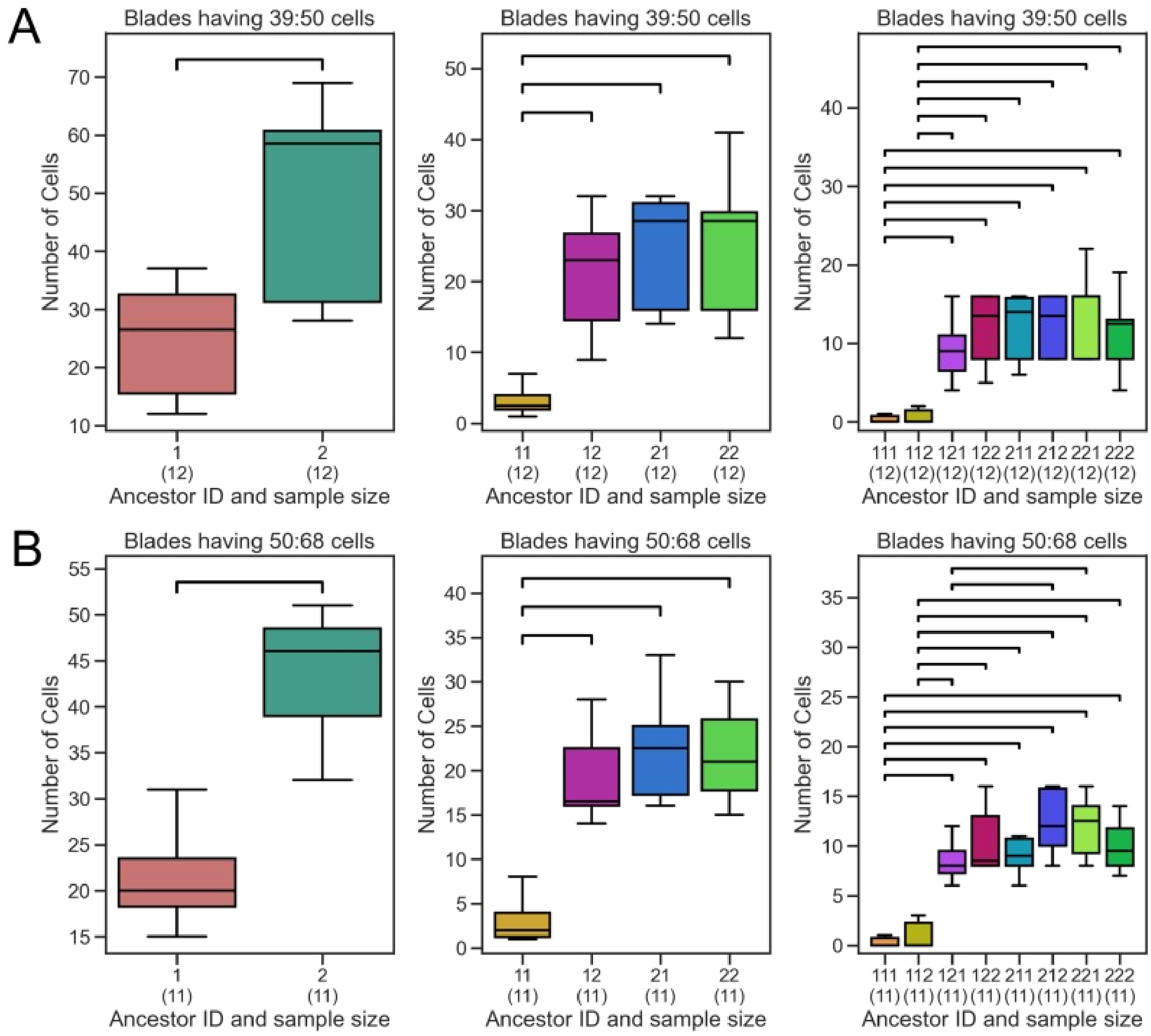


**Figure S3. Quantification of cellular lineage in the early embryo.**

The box plots illustrate the number of cells resulting from the mitotic activities of each embryo cell (named "ancestor cell") at the 2-cell (Anc1), 4-cell (Anc2) and 8-cell (Anc3) stages in embryos with 39-50-cells (A) and 50-68 cells (B). Statistics show that the mitotic activity of the basal cells is significantly lower than that of the other cells at the two developmental stages (Phase II) (Mann-Whitney U test, segments join samples with a p-value $< 0.05$). The identity of the "ancestor" cells is indicated under the boxes; the number of embryos is shown in parentheses.

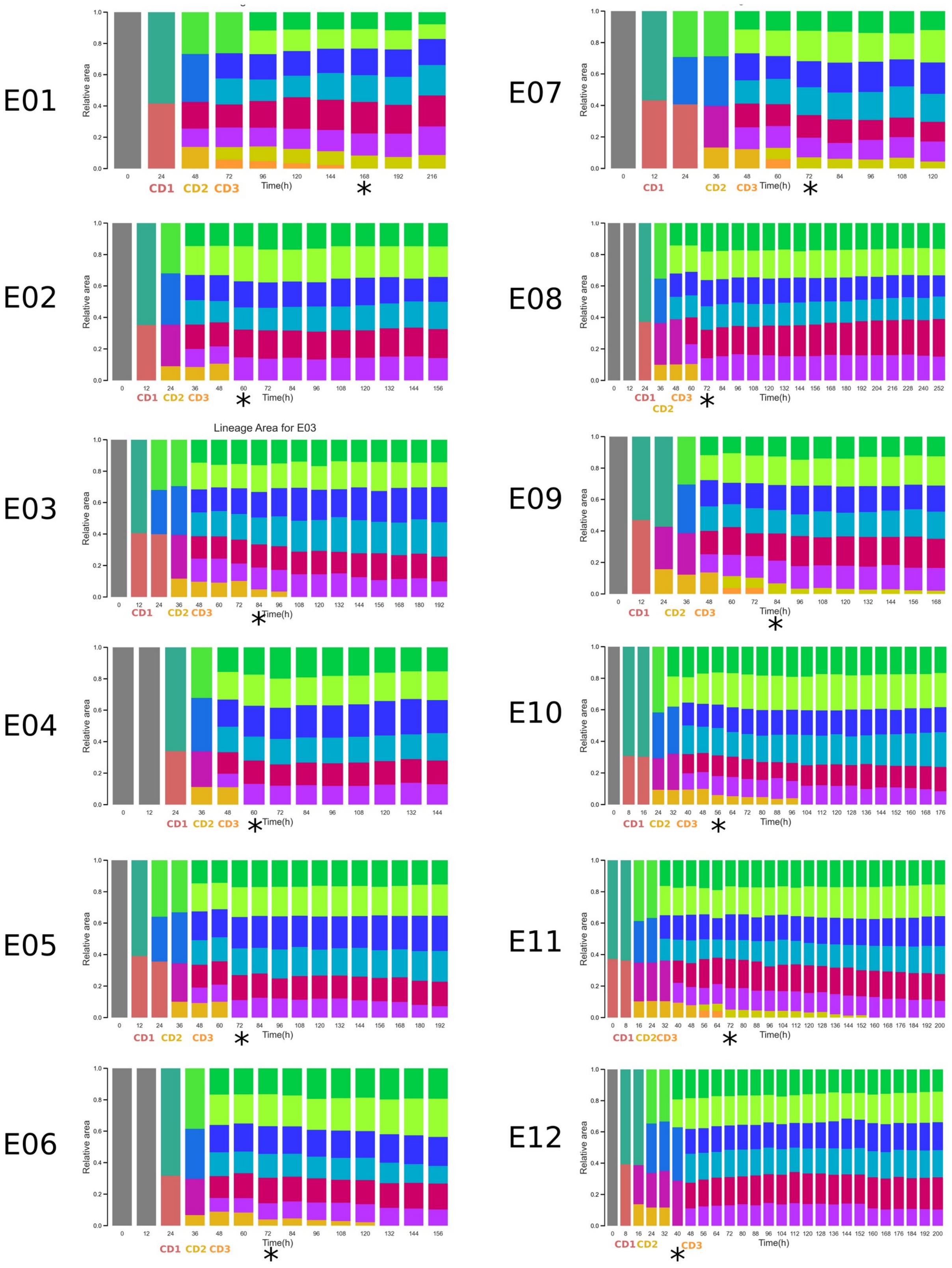


**Figure S4. Cell size and cell sectors resulting from cell divisions in the *Saccharina* embryos.**

Each plot corresponds to one embryo monitored for up to 252 hours (x-axis). Embryo E04 is also shown in Fig. 1E. Each column corresponds to a time point. Each colored box corresponds to one cell ("cold" colors for cells in the apical region and "warm" colors for cells in the basal region). The embryo develops from the zygotic stage (gray column on the left), up to 128 cells (last column on the right). The height of each colored sector corresponds to the surface area of the cells or their progeny relative to the whole embryo size. Only cells resulting from the first three rounds of cell division are given a distinct colored boxes. After that stage, each sector aggregates the progeny of one of the initial eight cells. CD# indicates the cell division round in the basal cell. Cells that differentiate into rhizoids disappear from the columns because they are no longer part of the embryo lamina. Asterisks indicate the time of the first rhizoid emergence.

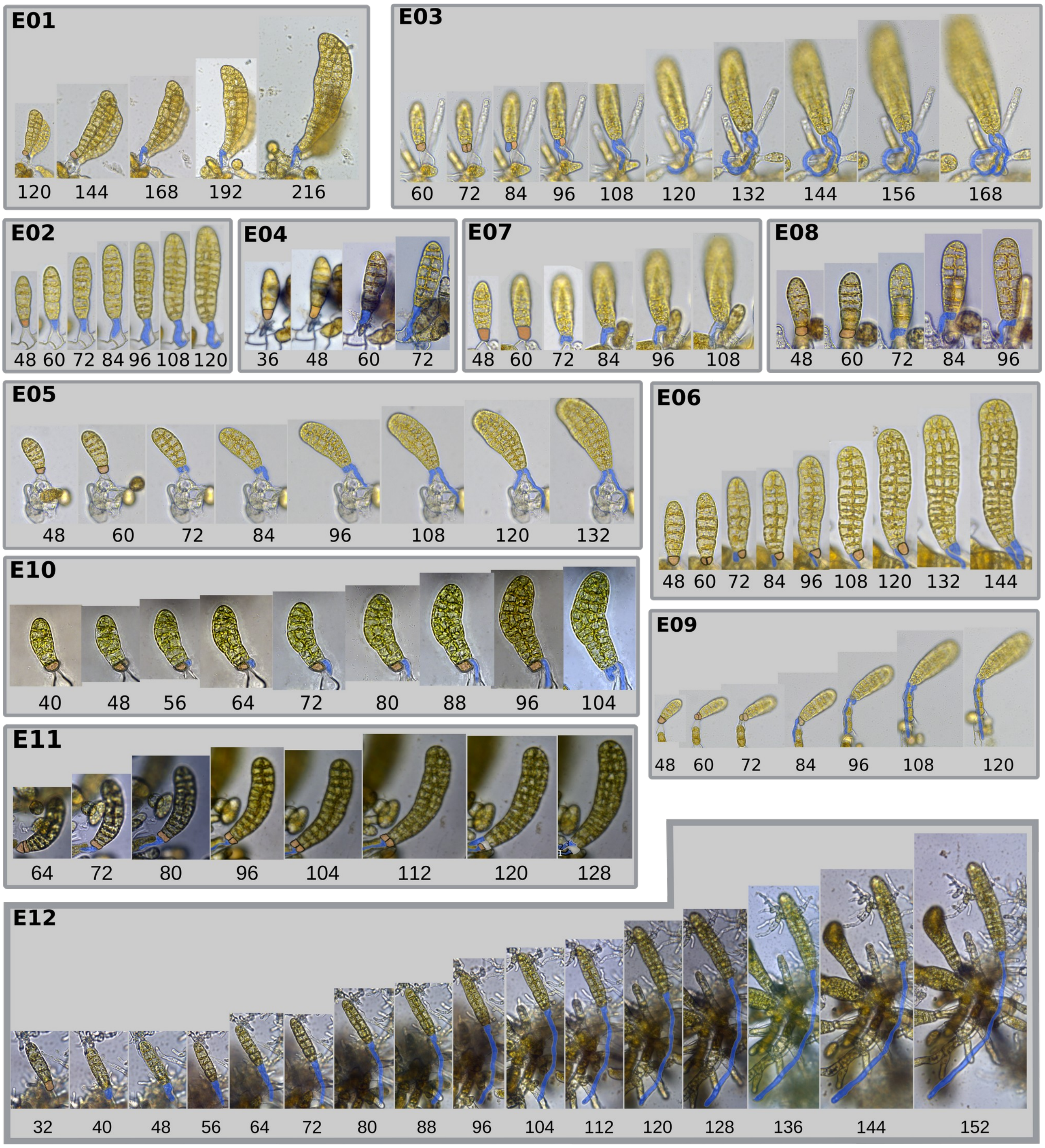


**Figure S5. Differentiation of basal cells into rhizoids observed in *Saccharina* embryos.**

The rhizoid initial, which is the basal cell about to differentiate into a rhizoid, is colored orange and the growing rhizoid is colored blue.

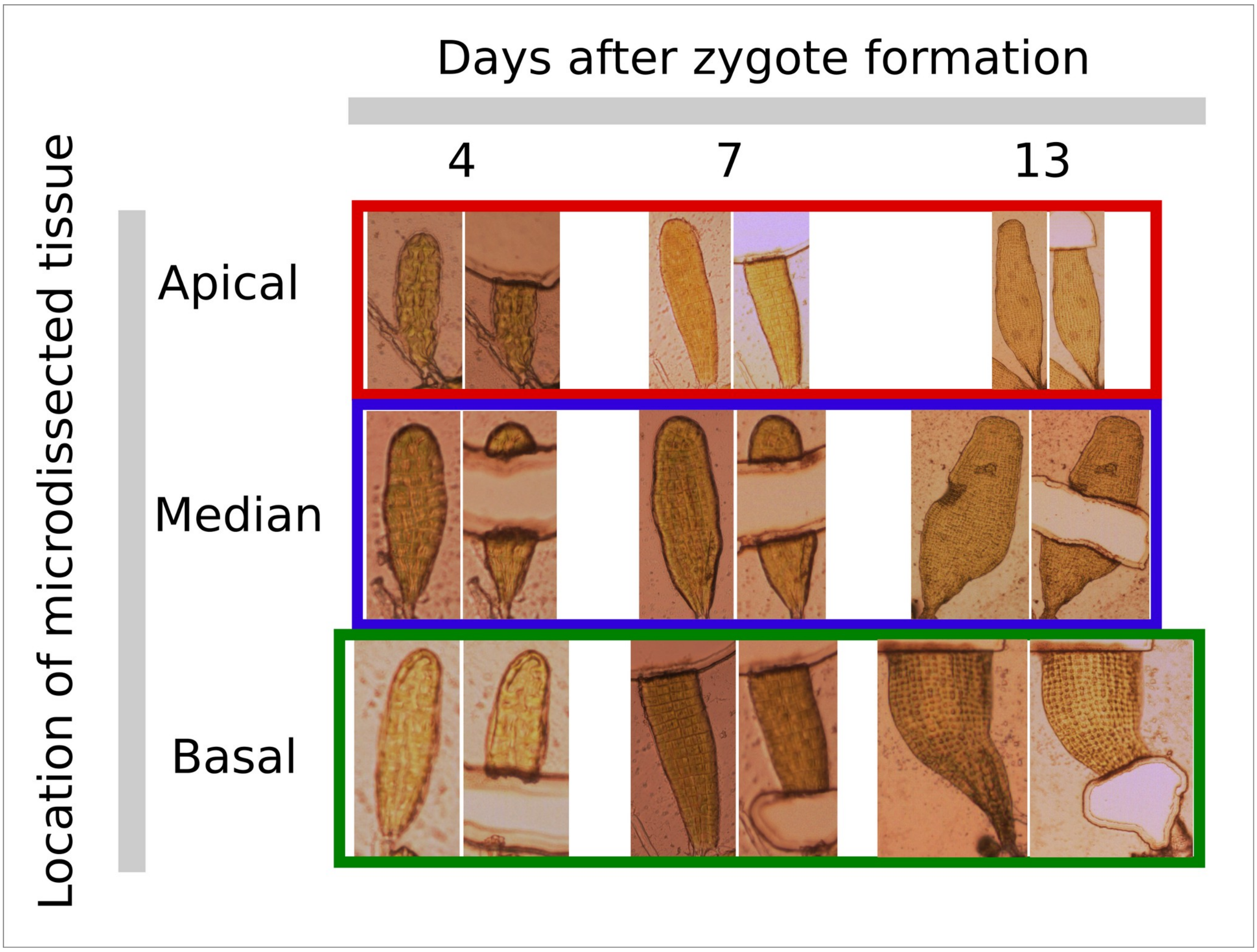


**Figure S6. Time and location of the ablated tissues in Phase II embryos prior to RNA-seq.** Individual cells or tissue sectors were microdissected using laser ablation before RNAs were extracted and sequenced (see Methods section for details of the experimental procedure).

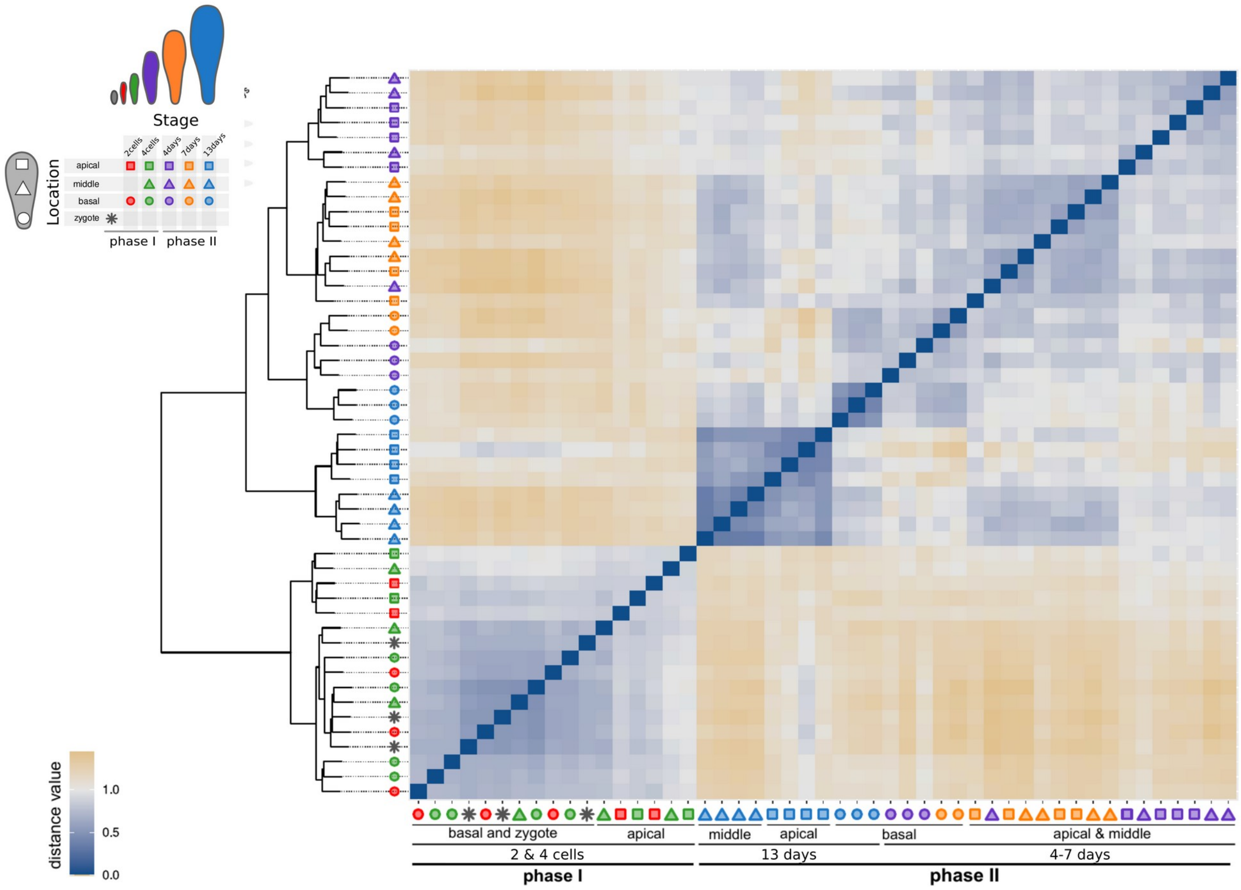


**Figure S7. Spearman correlation.**

This heatmap shows the distance between replicates based on Spearman's rank correlation coefficients. The blue-yellow color scale represents similar and dissimilar samples, respectively. A cladogram resulting from hierarchical clustering of the Spearman's rank correlations using Ward's minimum distance method is shown on the left. The same color and shape codes are used to represent the different microdissected samples as in Fig. 3B.

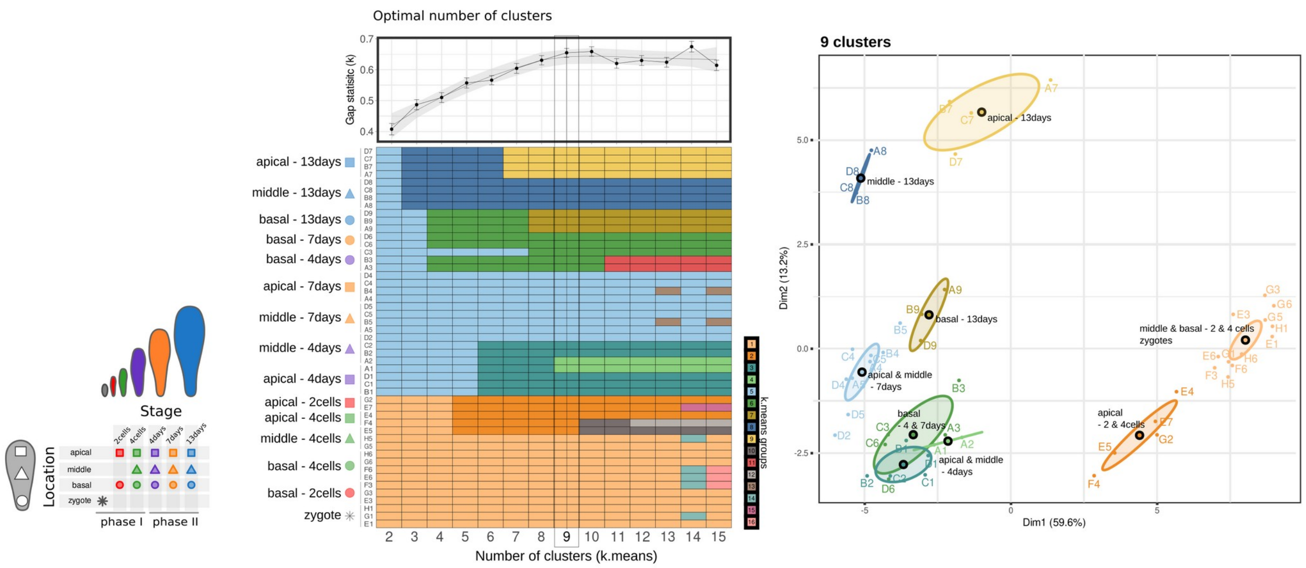


**Figure S8. K-means sequential clustering.**

K-means clustering of the replicates based on Spearman correlation distances. Left: K-means clustering for a series of 2-15 groups. The upper panel shows the gap statistic value, which estimates the optimal number of groups. The dotted line shows the local polynomial regression (less smooth), and the gray area shows the standard error. The lower panel shows a heatmap of cluster affiliation for each replicate. Right: K-means clustering of nine groups in a two-dimensional space using principal components. Small dots represent individual replicates. The large, black, circled dot is the cluster centroid. The ellipse represents cluster "confidence" and is filled with a color corresponding to the clusters represented in the left panel. The symbols are represented with the same color and shape codes as in Fig. 3B.

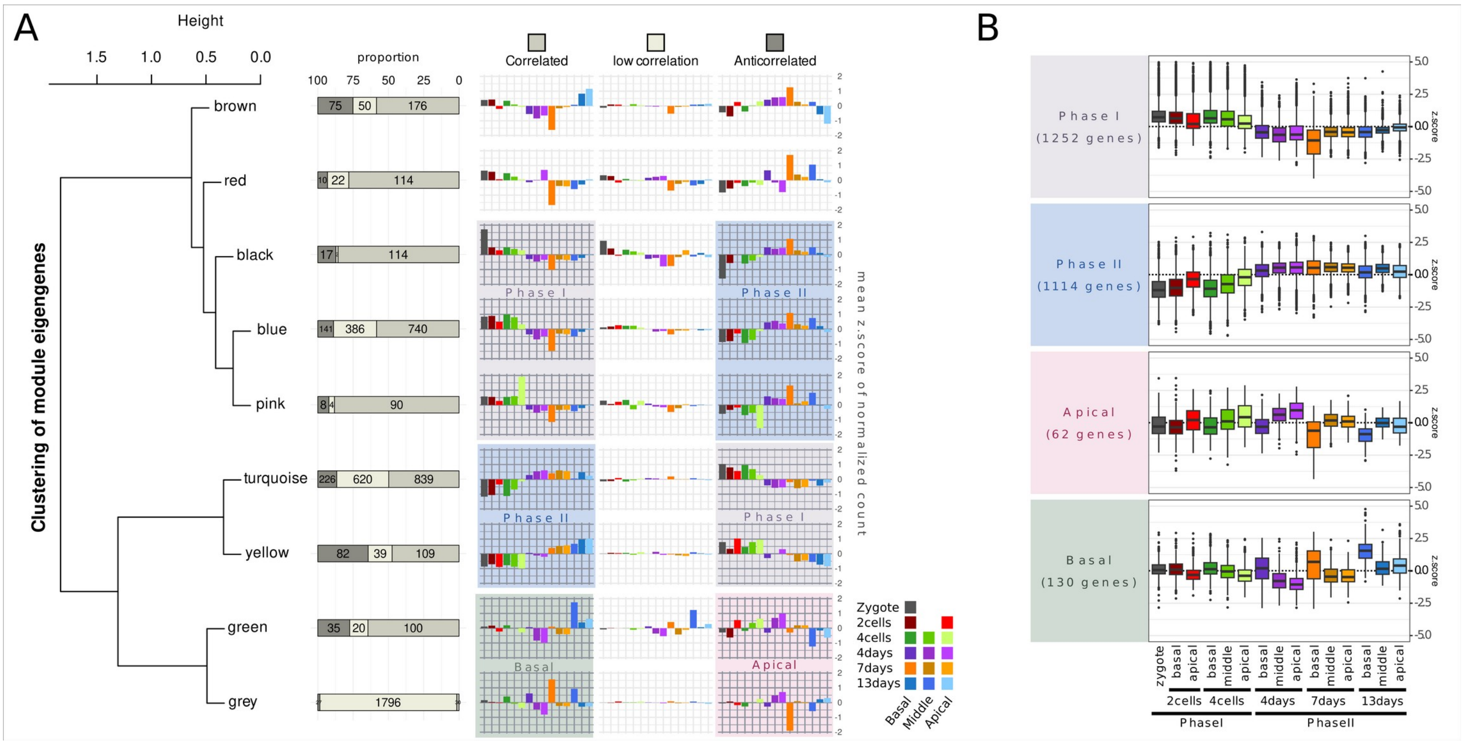


**Figure S9. Construction of groups of genes based on their expression profiles.**

Modules of co-expressed genes led to the identification of four overlapping groups of genes preferentially expressed in 1) Phase I, 2) Phase II, 3) Apical or 4) Basal cells/sectors of the embryo. **A.** The WGCNA gene co-expression analysis identified nine modules. Left panel: hierarchical clustering of the module eigengenes. Middle panel: horizontal barplot reporting the proportion of genes within each module that show "correlation" (weighted Pearson correlation ≥0.5; medium gray; highest number), "anticorrelation" (weighted Pearson correlation ≤0.5; dark gray) or "weak correlation" (-0.5≤ weighted Pearson correlation ≤0.5; light gray) with the module eigengene. The numbers correspond to the number of genes of each group. Right panel: barplot summarizing the gene expression profiles based on the mean z.scores of the normalized count. Positive z.scores show an over-expression level compared to the overall mean, negative z.scores show a lower expression level compared to the overall mean. **B.** Based on modules of co-expressed genes, we defined four groups of genes preferentially expressed in 1) Phase I (gray background), 2) Phase II (blue background), 3) Apical (pink background) or 4) Basal (green background) cells/sectors of the embryo. Box plots show the distribution of z.scores of the normalized counts per condition (stage and location) for each of the four groups.

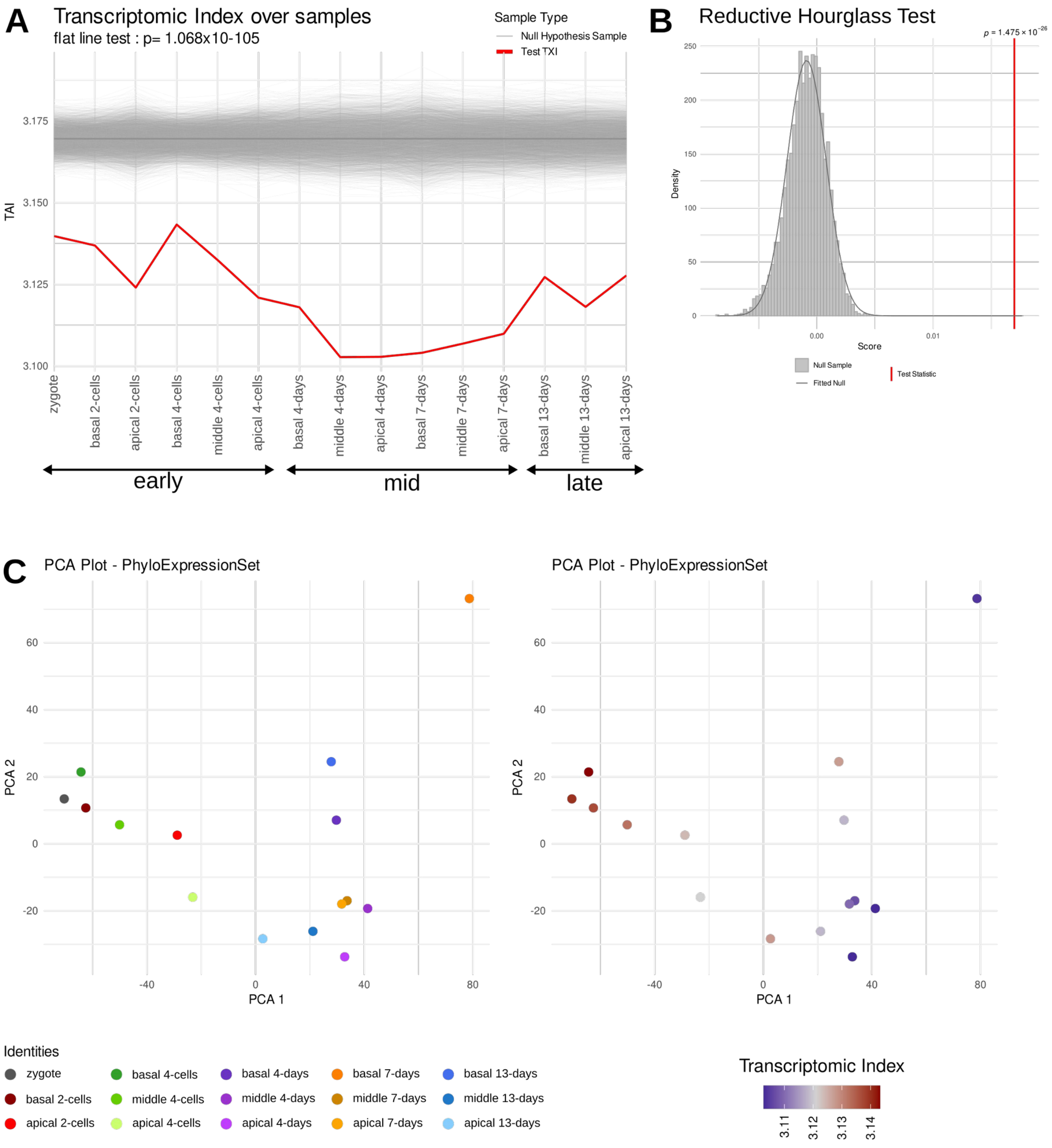


**Figure S10. Transcriptome age analysis using phylostratigraphy**

Transcriptomic Age Index (TAI) represents the overall age of a transcriptome based on the age and expression level of individual genes. The higher the TAI, the younger the transcriptome (recent genes are expressed at a higher level). The lower the TAI, the older the transcriptome (ancestral genes are expressed at a higher level).

**A.** Flatline test evaluating whether the observed TAI profile (red line) is significantly different from a flatline. The subtitle indicates the associated p-value. The light gray lines represent individual permutations of null hypothesis (flat line) and the horizontal gray line shows the mean of those null distributions. **B.** Dimensional reduction plot using the PCA method to represent the relative positioning of samples according to gene expression. The right panel colors samples by identity, and the left panel colors samples by TAI values. **C.** Reductive Hourglass Test, evaluating whether the observed TAI profile (red line form panel A) significantly deviates from the null hypothesis towards an hourglass pattern. The TAI values are lower in the mid-stage than in the early and late stages. We considered the “zygote”, “2cells and “4cells” samples to be early-stage, “4days” and “7days” samples to be mid-stage and “13days” samples to be “late-stage”. The gray bars and line represent the score values and distribution, respectively, of the individuals permutations of the null hypothesis (not hourglass); the vertical red line represents the score value for the observed data with the associated p-value indicated on the line.

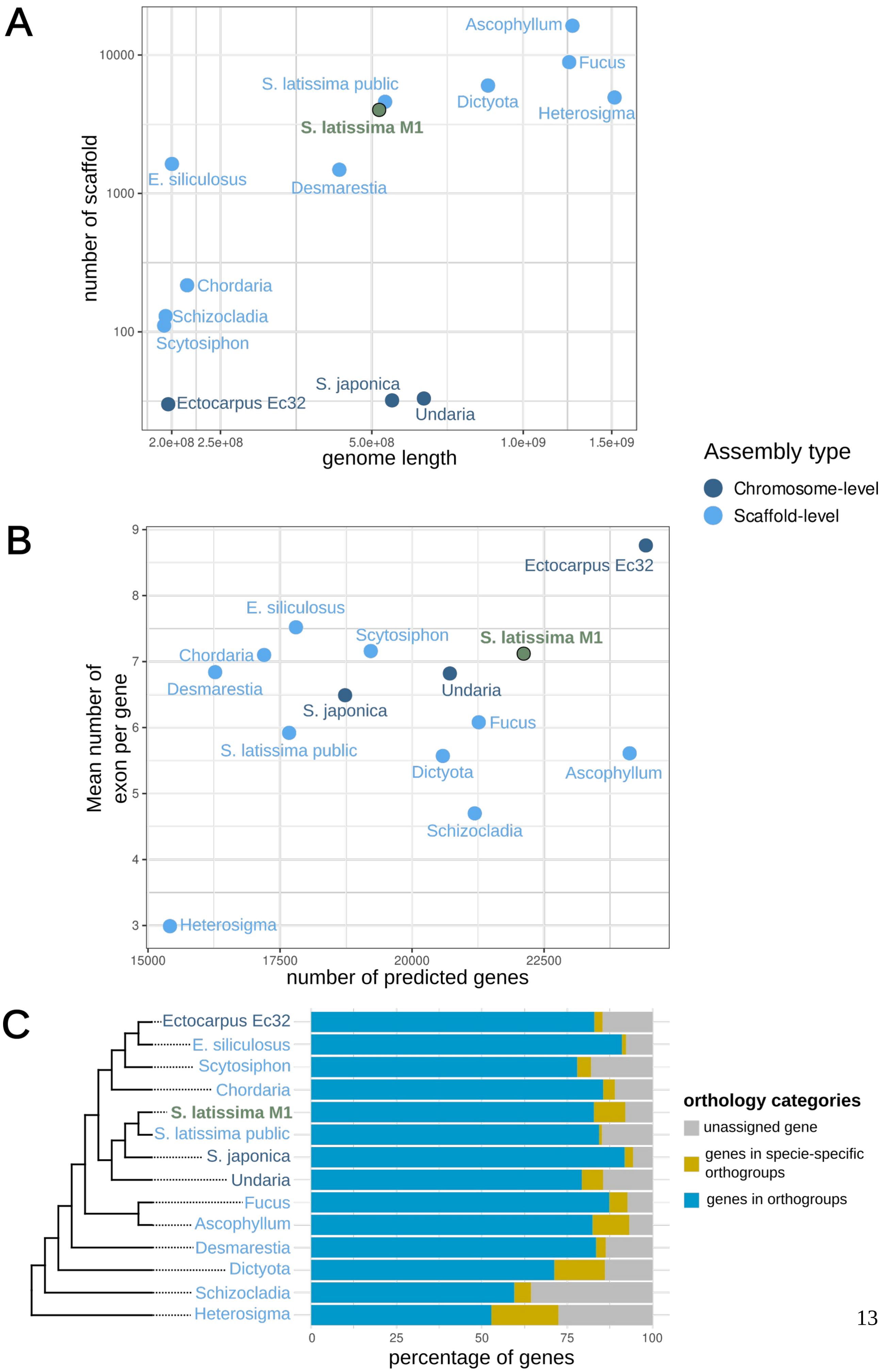
A
number of scaffold
genome length
10000
1000
100
2.0e+08 2.5e+08
5.0e+08
1.0e+09
1.5e+09
Ascophyllum
Fucus
S. latissima public
Dictyota
Heterosigma
S. latissima M1
E. siliculosus
Desmarestia
Chordaria
Schizocladia
Scytosiphon
S. japonica
Ectocarpus Ec32
Undaria
Assembly type
Chromosome-level
Scaffold-level
B
Mean number of exon per gene
number of predicted genes
9
8
7
6
5
4
3
15000
17500
20000
22500
Ectocarpus Ec32
E. siliculosus
Scytosiphon
S. latissima M1
Chordaria
Desmarestia
Undaria
S. japonica
Fucus
S. latissima public
Dictyota
Ascophyllum
Schizocladia
Heterosigma
C
Ectocarpus Ec32
E. siliculosus
Scytosiphon
Chordaria
S. latissima M1
S. latissima public
S. japonica
Undaria
Fucus
Ascophyllum
Desmarestia
Dictyota
Schizocladia
Heterosigma
0
25
50
75
100
percentage of genes
orthology categories
unassigned gene
genes in specie-specific orthogroups
genes in orthogroups

**Figure S11: Genome sequence quality**

The quality metrics of the newly assembled *S. latissima* M1 genome are compared to those of a selection of current, publicly available genomes. The considered reference species are accessible in the Table S5. **A.** Genome fragmentation: dot-plot representing total genome length against the total number of scaffolds in the assembly. **B.** Gene model fragmentation: dot-plot representing the total number of predicted genes against the mean number of exons per gene. **C.** Gene prediction orthology (orthofinder results). Left panel: cladogram based on the phylogenetic relationship of the considered species. Right panel: bar plot showing the proportion of predicted genes included in an orthology relationship with other species.
